# Hindbrain explants enable multimodal and longitudinal analysis of the developing olivo-cerebellar circuit at single-cell resolution

**DOI:** 10.64898/2026.08.26.747061

**Authors:** Elena Baz-Badillo, Charlotte Taeger, Margaux Saint-Martin, Charles Ducrot, Lucie Franco, Thaïs Verschaeve, Alexandre Favereaux, Elena Avignone, Mathieu Letellier

## Abstract

Experimental models that preserve native mammalian CNS circuitry while enabling longitudinal analysis of circuit assembly at single-cell resolution remain scarce, limiting mechanistic studies and therapeutic discovery. Here, we establish embryonic mouse hindbrain explants as a scalable *in vitro* model that maintains the long-range olivo-cerebellar circuit while providing direct experimental access to both pre- and postsynaptic neurons. The preparation supports repeated live imaging, targeted single-cell manipulation and labelling, electrophysiology, ultrastructural analysis, and single-cell RNA sequencing during circuit assembly. Hindbrain explants faithfully recapitulate key features of olivo-cerebellar organization and development, including cytoarchitecture, synaptic organization and maturation, neuronal differentiation, and spontaneous network activity while preserving developmental glial features. By combining developmental and physiological fidelity with longitudinal multimodal accessibility, this resource bridges the gap between reductionist cultures and technically demanding *in vivo* approaches, providing a versatile and ethical model for investigating the molecular and cellular mechanisms of cerebellar circuit assembly and disease.

## Introduction

Neural circuit assembly requires the coordinated differentiation of multiple neuronal populations, the establishment of long-range connectivity and the progressive refinement of synaptic interactions into mature functional networks. Dissecting these processes requires experimental models that preserve native circuit architecture while enabling longitudinal observation and manipulation at single-cell resolution ^1^. Although in vivo approaches have considerably advanced our understanding of neural development, experimental access to intact central mammalian circuits during the critical periods of synapse formation and refinement remains limited.

The olivo-cerebellar circuit provides one of the best-established models for investigating developmental circuit assembly. Purkinje cells (PCs), the sole output neurons of the cerebellar cortex, receive two anatomically and functionally distinct excitatory inputs: the climbing fibers (CFs) arising from inferior olivary neurons (IONs), and the parallel fiber (PFs) from cerebellar granule cells (GCs). CFs establish hundreds of synapses onto proximal PC dendrites and undergo a remarkable developmental refinement in which multiple competing inputs are progressively eliminated, leaving most PCs innervated by a single dominant CF in the mature cerebellum ^2–5^. Mature CF-PC synapses evoke large all-or-none complex spikes (CSs), providing a unique physiological signature that directly reports functional olivo-cerebellar connectivity ^3,6,7^. In parallel, GCs receive information from pontine, vestibular and spinocerebellar pathways through mossy fibers (MFs) and establish PF synapses onto distal PC dendrites ^8–11^. The coordinated maturation of these excitatory inputs, together with inhibitory interneurons and glial populations, generates the highly organized cerebellar circuitry underlying motor coordination, timing and learning ^12^, and increasingly recognized for its roles in higher-order functions, including emotional processing, social behaviour, language and sleep as well as in neurodevelopmental disorders such as autism spectrum disorders ^13–18^.

Despite its defined anatomical and functional properties, the olivo-cerebellar circuit remains difficult to investigate as an integrated system. While the cerebellar cortex is relatively accessible in vivo, the inferior olive is deeply embedded within the ventral brainstem, limiting direct investigation. Moreover, many of the defining events of circuit assembly, including axon targeting, synaptogenesis, dendritic growth and CF refinement, occur during embryonic and early postnatal development, when chronic imaging, targeted manipulations and electrophysiological recordings remain particularly challenging.

Current *in vitro* models only partially overcome these limitations. Dissociated cultures provide excellent experimental accessibility but sacrifice native cytoarchitecture, long-range connectivity and physiological activity ^19–23^, whereas organotypic slices preserve local tissue organization but disconnect PCs from their brainstem afferents and downstream targets, altering neuronal survival, synaptic maturation and circuit refinement ^24–27^. Heterochronic co-cultures of postnatal cerebellar slices with embryonic brainstem tissue partially restore CF innervation but necessarily disrupt developmental timing, synaptic refinement and topographic organization while introducing slicing-induced inflammatory responses that may influence neuron-glia interactions ^28–30^. Consequently, no preparation currently offers both preservation of intact long-range olivo-cerebellar connectivity and the broad experimental accessibility required to investigate circuit assembly across molecular, structural and functional scales.

Hindbrain explant cultures offer a unique solution to this challenge. Prepared from mouse or chick embryos, these unsliced “open-book” preparations preserve the cerebellum and brainstem nuclei connected, including inferior olive, while providing direct access to both pre- and postsynaptic compartments. Explants can be maintained throughout the entire period of olivo-cerebellar circuit assembly, from initial axon targeting to mature synaptic refinement **(Sup. Fig. 1A)**. While two studies have highlighted the utility of this system for investigating early cerebellar circuit development and refinement in chick or mouse ^31,32^ it remains incompletely characterized, limiting its broader use. In particular, it is still unknown whether neurons in hindbrain explants retain their molecular identity and connectivity, recapitulate normal developmental trajectory and whether these cultures develop glial reactivity as usually observed in primary or slice cultures.

**Figure 1:**
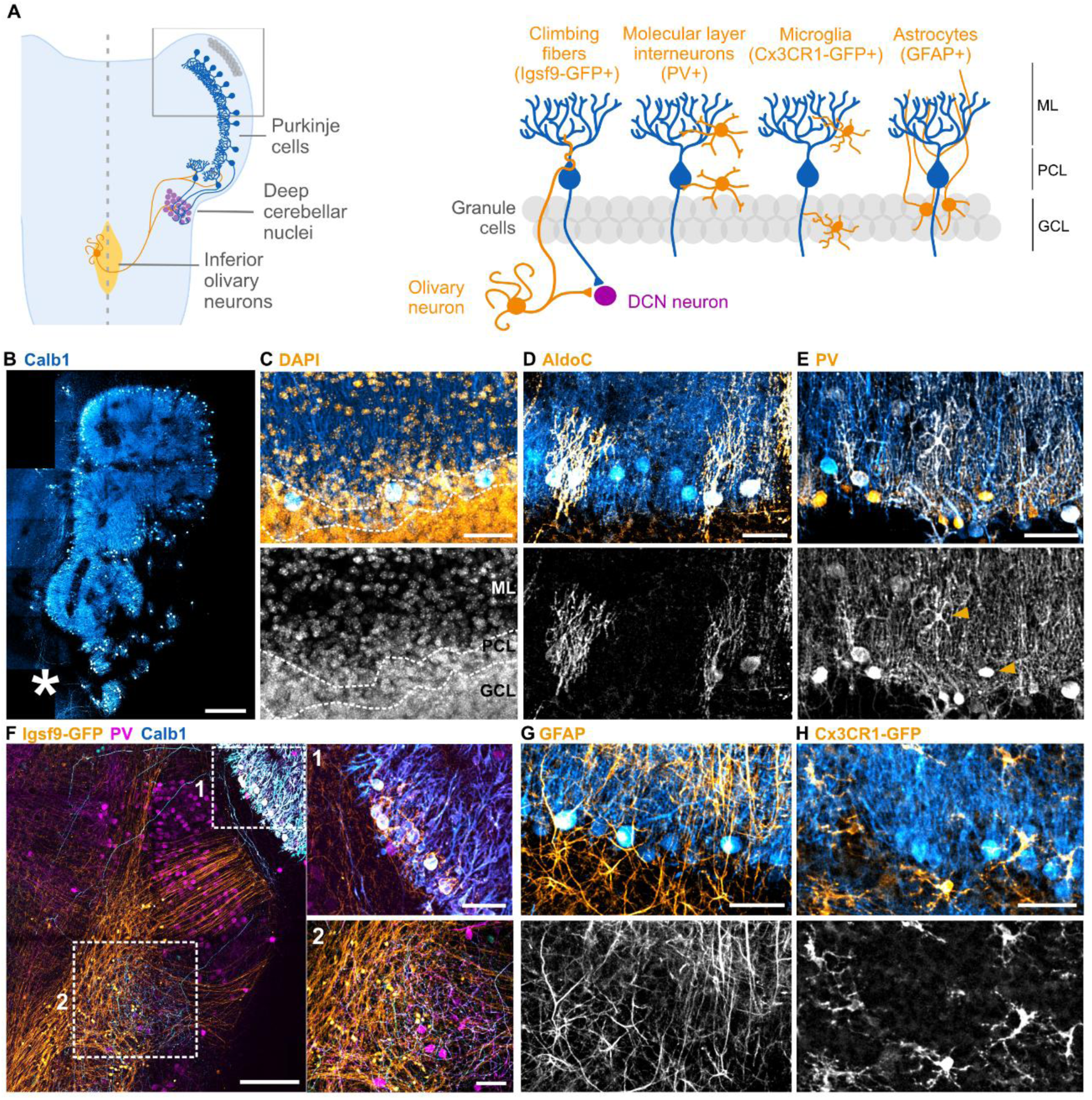
Hindbrain explants recapitulate the cellular organization of the cerebellar cortex. All confocal images correspond to maximum intensity projections (MIP) of image stacks, with PCs immunolabelled for Calb1 in blue. **(A)** Scheme showing the position of IONs, DCN and cerebellar cortex in explant (right), and the organization of cerebellar cortex in explants (left). **(B)** Low magnification image showing the distribution of Calb1+ PCs in the cerebellar plate. Scale bar = 300 μm. The asterisk indicates the position of Calb1+ PC axon terminals. **(C)** Top: Organization of Calb1+ PCs (blue) relative to GCL (DAPI, orange). Bottom: image showing DAPI staining only. Dashed lines delimit the ML, PCL and GCL. **(D)** Top: image showing Calb1+ (blue) and Calb1+/AldoC+ PCs (orange) in the cerebellar plate. Bottom: AldoC immunolabelling only. Median percentage (IQR) of AldoC+ PCs = 50.2% (8.53%) (n = 5 cerebellar plates from 3 explants, 180-290 PCs analyzed per cerebellar plate) **(E)** Top: distribution of PV+ interneurons (orange) relatively to PCL and ML (Calb1 immunolabelling, blue). Bottom: PV immunolabelling only. Arrowheads indicate PV+/Calb1-cells. **(F)** Left: position of DCN posterior to the cerebellar plate in a Igsf9-GFP explant (to visualize CFs, in orange) immunostained for Calb1 (blue) and PV (magenta). Asterisk in (B) indicates the approximate position in the explant. Scale bar = 150 μm. Right, top: high magnification of the area framed with dashed line, showing GFP+ CFs (orange) innervating Calb1+ PCs (blue) in the cerebellar plate. Right, bottom: high magnification of the area framed with solid line showing DCN containing PV+ neurons (magenta) and axon terminals from both GFP+ CFs (orange) and Calb1+ PCs (blue). Scale bars = 50 μm. **(G)** Top: morphology and distribution of astrocytes immunolabelled for GFAP (orange) relative to Calb1+ PCs (blue). Bottom: GFAP immunostaining only. **(H)** Top: morphology and distribution of GFP+ microglia from Cx3cr1-GFP explant (orange) relative to Calb1+ PCs (blue). Bottom: GFP+ microglia only. **(C, D, E, G, H)** Scale bar = 50 μm.

Here, we establish embryonic mouse hindbrain explants as a multimodal system for investigating cerebellar circuit development at single-cell resolution. We demonstrate that explants faithfully reproduce the structural organization, developmental progression, functional maturation and transcriptional programs of the olivo-cerebellar circuit while providing direct experimental access to both pre- and postsynaptic neurons. By combining live imaging, targeted single-cell manipulation, electrophysiology, immunohistochemistry and Patch-seq, this resource enables integrated investigation of neural circuit assembly across molecular, cellular and functional scales. Together, our work positions hindbrain explants as a versatile platform for longitudinal investigation of cerebellar development, neuron-glia interactions and mechanisms underlying neurodevelopmental disease at cellular resolution.

## Results

### Hindbrain explants self-organize into mature cerebellar architecture

We first assessed the extent to which cerebellar cytoarchitecture is preserved in explants prepared from E14.5 mouse embryos **(Sup. Fig. 1A-1F)** and cultured for 28 days (DIV28), i.e., equivalent to postnatal day 21 (P21). We examined the presence of major neuronal and glial cell types and their organization into the characteristic layers of the mature cerebellum: the molecular layer (ML), PC layer (PCL), and GC layer (GCL). Explants were prepared from Igsf9-eGFP or Cx3CR1-GFP embryos, in which IONs or microglia, respectively, express GFP ^33,34^. They were subsequently immunostained for Calbindin-1 (Calb1) and Aldolase C (AldoC) to label PCs, Parvalbumin (PV) to label interneurons and GFAP to label astrocytes, with DAPI marking nuclei **(Fig. 1A)**.

Calb1-positive (Calb1+) PCs displayed aligned cell bodies and typical polarized dendritic trees organizing the PCL and the ML, respectively **(Fig. 1B-C)**. A dense population of DAPI-positive nuclei, presumably GCs, was located beneath the PCL, defining the GCL **(Fig. 1C)**. Approximately 50% of PCs were also immunopositive for aldolase C / zebrin II, indicating that PC molecular heterogeneity is preserved in the explants **(Fig. 1D)**. Zebrin II-positive PCs were arranged in patches in the cerebellar plate, similar to the zebrin bands configuration *in vivo* ^35^ (**Sup. Fig. 2**). PV+, Calb1-neurons were present in the ML, as well as within PC and GC layers, suggesting the presence of molecular interneurons (MLI) and Golgi cells (GoC) **(Fig. 1E)**. Immediately posterior to the cerebellar plate, a population of PV+/Calb1-neurons was observed, targeted by both PCs axon terminals as well as CF terminals from GFP+ IONs, indicating the position of deep cerebellar nuclei (DCNs) **(Fig. 1F)**.

**Figure 2:**
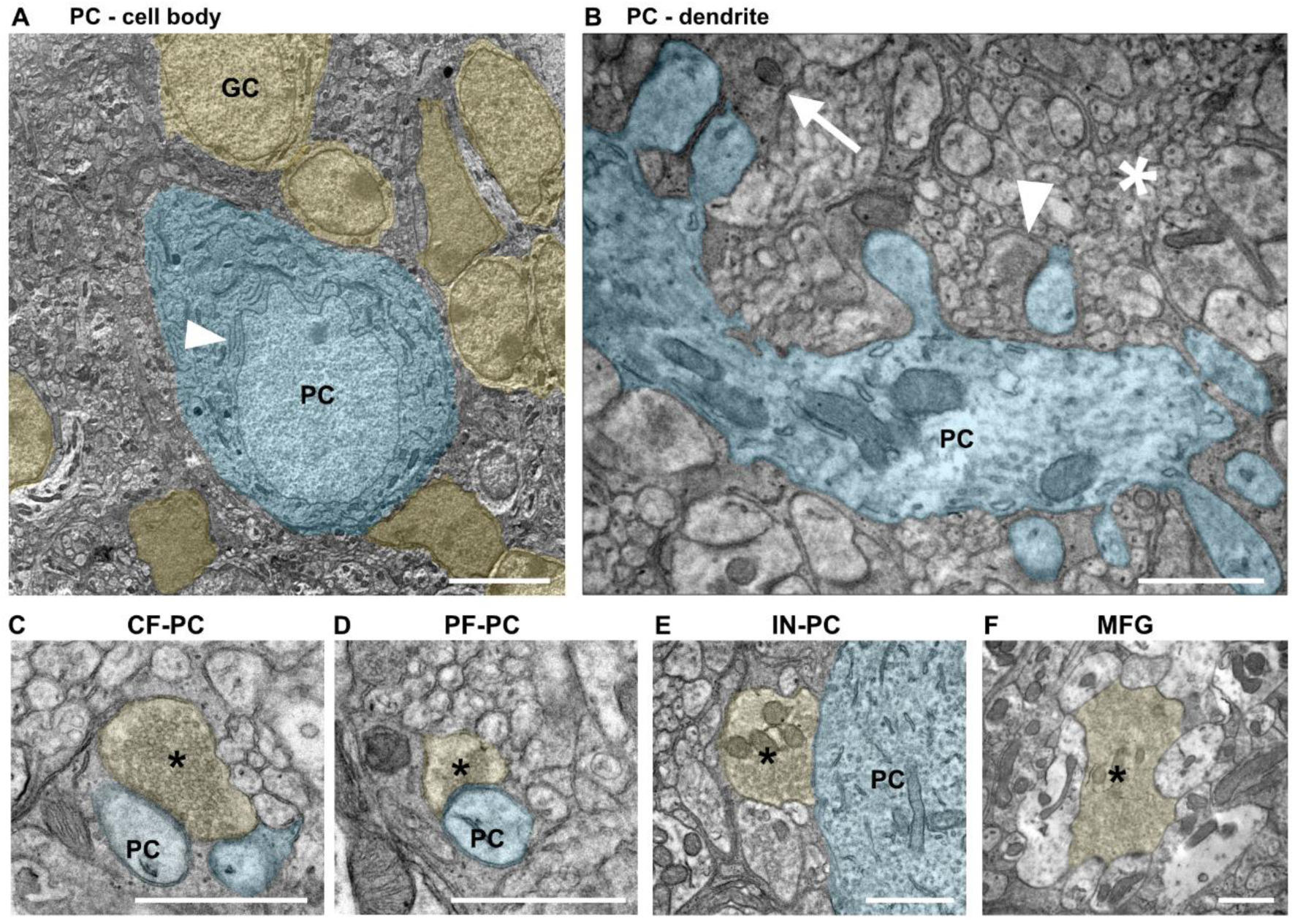
Hindbrain explants develop mature cerebellar cellular and synaptic ultrastructure. **(A)** TEM image showing a PC cell body (blue) in a DIV30 explant. Arrowhead indicates Nissl bodies (stacked endoplasmic reticulum). Asterisk indicates the position of GCL (yellow). Scale bar = 5 μm. **(B)** TEM image showing a PC dendrite (blue) in the ML. Arrowhead indicates a putative PF terminal. Arrow indicates a putative CF terminal. Asterisk indicates PF axons. **(C)** TEM image showing a putative CF synapse on PC dendritic spine. **(D)** Putative PF synapse on PC dendritic spine. **(E)** Putative inhibitory synapse on PC cell body. **(F)** MFG within the GCL. **(C-F)** Scale bar = 1 μm.

GFAP+ cells adjacent to PCs somata extended radial processes throughout the ML, ensheathing PC dendrites and forming a scaffold-like structure reminiscent of Bergmann glia *in vivo* **(Fig. 1G)**. Cx3CR1-GFP+ microglia were observed throughout the different cerebellar layers and displayed ramified morphology comparable to *in vivo* condition ^36^ **(Fig. 1H)**.

Overall, the preservation of cerebellar layering and major neuronal and glial populations indicates that hindbrain explants retain the general cellular organization of the mature cerebellum.

### Long-term maintenance of cerebellar ultrastructure in hindbrain explants

We next asked whether this preserved cellular organization extends to the ultrastructure of cerebellar neurons and synapses, a prerequisite for normal circuit function ^37–42^. To this end, DIV30 explants were processed for transmission electron microscopy (TEM) **(Fig. 2A-F**). PC cell bodies were readily identified by their drop-shaped morphology, their large euchromatic nucleus containing a marked nucleolus and cytoplasm containing highly characteristic stacked rough endoplasmic reticulum cisternae (Nissl bodies), a well-developed Golgi apparatus and numerous mitochondria **(Fig. 2A)**. Beneath the PC layer, we observed packed small cell bodies with dense heterochromatin that we identified as putative granule cells **(Fig. 2A)**.

In the ML, transverse sections of thin PF axons were abundant **(Fig. 2B)**. PC dendrites displayed numerous dendritic spines receiving asymmetric, excitatory synapses of distinct morphologies **(Fig. 2B)**. In agreement with the descriptions in the literature, electron-dense presynaptic terminals containing large, uniformly distributed vesicles apposed to dendritic spines emerging from thick dendrites in proximity of PC somata were identified as CF boutons **(Fig. 2C)**. On the other hand, lighter boutons in which vesicles accumulated near the active zone were consistent with PF synapses **(Fig. 2D)**. In the PC cell body, terminals with pleomorphic vesicles formed symmetric synapses, identified as inhibitory contacts **(Fig. 2E)**. Within the GCL, mossy fiber glomeruli (MFG) were readily recognized by their characteristic rosette-like organization and convergence of morphologically distinct boutons **(Fig. 2F)**. No signs of degeneration, autophagy or cellular atrophy were observed.

Together, these observations demonstrate that DIV30 hindbrain explants maintain the characteristic ultrastructure of the intact cerebellum *in vivo*, indicating that neuronal morphology and synaptic architecture are maintained in long-term culture conditions.

### Hindbrain explants recapitulate spatio-temporal development of Purkinje cell synaptic connectivity

To determine whether the cellular and synaptic organization observed in mature explants emerges through the developmental sequence described *in vivo* ^42,43^, we examined the spatiotemporal maturation of excitatory and inhibitory inputs onto PCs. Igsf9-GFP explants, in which IONs express GFP, were analyzed at DIV7-8, DIV14, DIV21 and DIV26, corresponding approximately to P0-P1, P7, P14 and P19, respectively. PCs were identified by Calb1 immunolabelling, while CF terminals were unambiguously identified by combined GFP expression and immunoreactivity for the vesicular glutamate transporter VGLUT2. PF terminals were immunolabelled for the vesicular glutamate transporter VGLUT1, with VGLUT2 additionally marking immature PF terminals. Inhibitory terminals arising from recurrent PC collaterals, MLIs and GoCs were identified by immunolabelling the vesicular GABA transporter VGAT.

At DIV7–8 (∼P0), PCs displayed a fusiform morphology, with one or a few processes extending from the soma, closely resembling their *in vivo* counterparts at P0–P1 ^44^. CF terminals were already detected on PC somata, consistent with the “creeper stage” of CF innervation described *in vivo* ^42,45^ **(Fig. 3A-3B)**. In contrast, VGLUT1 immunoreactivity was largely absent from the cerebellar plate, indicating that PF innervation had not yet emerged **(Fig. 3A, 3J)**. Inhibitory terminals were already present within the PCL at a density comparable to that of CF terminals (VGLUT2+ puncta density: 0.0252±0.0095 puncta·µm^-2^; VGAT+ puncta density: 0.0264±0.0068 puncta·µm^-2^) **(Fig. 3B, 3K)**, consistent with the early establishment of inhibitory innervation *in vivo* ^42^.

**Figure 3:**
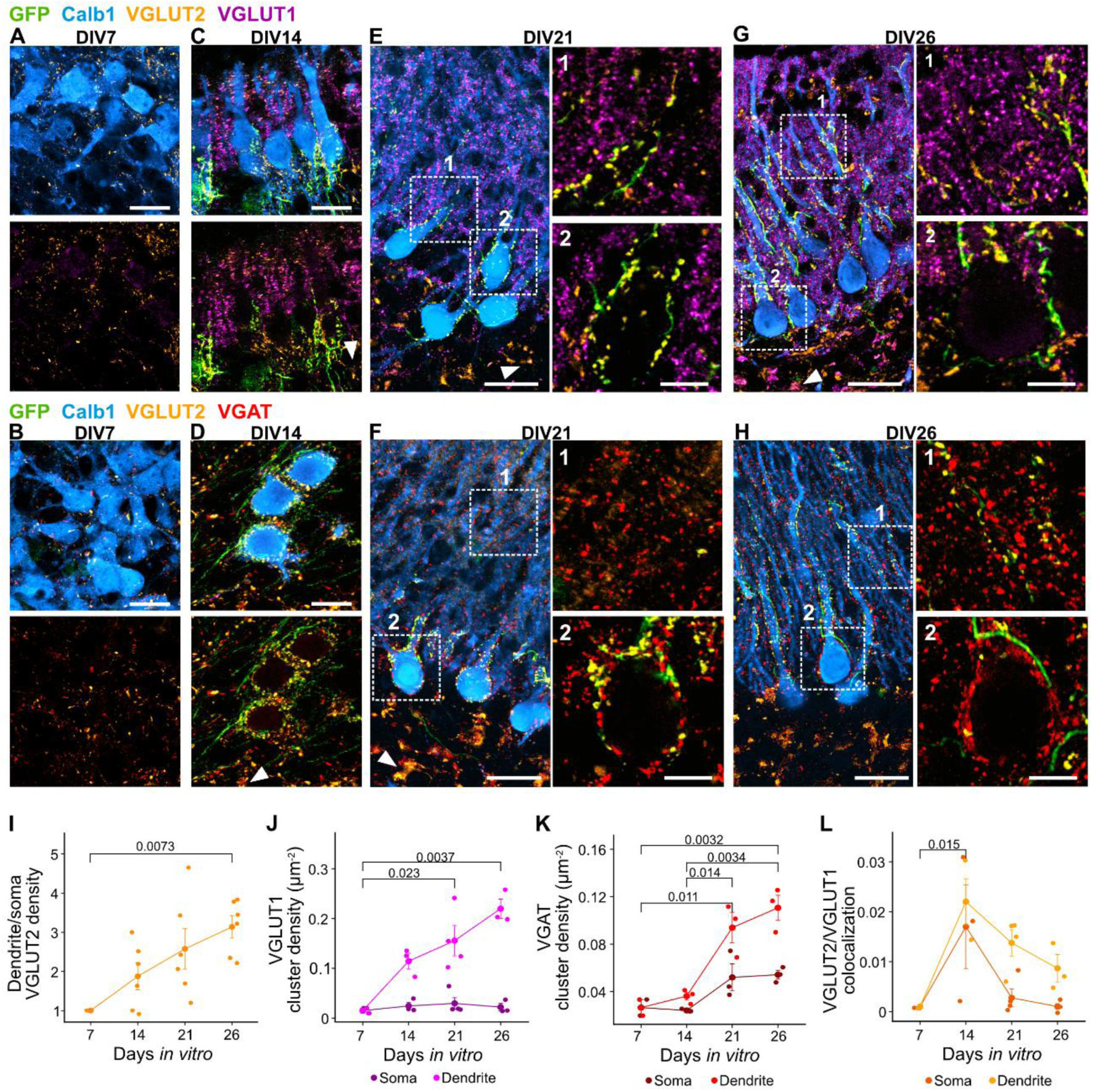
PC synaptic connectivity undergoes coordinated spatiotemporal maturation in hindbrain explants. **(A-H)** Maximum intensity projections of confocal images stacks in the cerebellar plates from Igsf9-GFP explants (10 μm). **(A, C, E, G)** DIV7 to DIV26 Igsf9-GFP explants containing GFP+ CFs (green) and immunostained for Calb1 (blue), VGLUT2 (yellow) and VGLUT1 (magenta). **(B, D, F, H)** DIV7 to DIV26 Igsf9-GFP explants containing GFP+CFs (green) and immunostained for Calb1 (blue), VGLUT2 (yellow) and VGAT (red). **(A-D)** Scale bar = 20 μm. **(E-H)** Low magnification images; scale bar = 30 μm. Zoomed images scale bar = 10 μm. **(C-G)** Arrowheads indicate MFG. **(C)** Arrow indicates VGLUT1/VGLUT2 coexpression. **(I)** Dendritic / somatic VGLUT2 puncta density ratio (DIV7 n = 4 explants; DIV14 n =7; DIV26 n = 7). ANOVA p = 0.008. Displayed p value corresponds to Tukey’s HSD pairwise comparisons. **(J)** VGLUT1 density (puncta·μm^2^) in soma (purple) and dendrites (magenta) (DIV7: n = 2 explants; DIV14: n = 3; DIV21: n = 4; DIV26: n = 3). ANOVA across time for dendrites: p = 0.005. Kruskal-Wallis for soma p = 0.67. **(K)** VGAT density (puncta·μm^2^) in soma (dark red) and dendrites (red) (DIV7 n = 2 explants; DIV14 n = 3; DIV21 n = 3; DIV26 n = 3). ANOVA across time for dendrites: p = 0.001. ANOVA across time for soma: p = 0.032. Tukey’s HSD for soma: p > 0.06 for all comparisons. **(L)** VGLUT2+/VGLUT1+ colocalization (puncta·μm^2^) in soma (dark orange) and dendrites (orange) (DIV7 n = 2 explants; DIV14 n = 3; DIV21 n = 4; DIV26 n = 4). ANOVA across time for dendrites: p = 0.018. Kruskal-Wallis for soma: p = 0.163. **(I-L)** Each data point corresponds to one explant. Data displayed corresponds to the mean±SEM per age. Displayed p values correspond to Tukey’s HSD across time for dendrites.

By DIV14 (∼P7), PCs showed clear signs of dendritic maturation. Some exhibited a stellate morphology with short, non-polarized dendrites, whereas others displayed an emerging primary dendrite **(Fig. 3C-3D)**, marking the onset of dendritic polarization. CF terminals formed characteristic “pericellular nests” around PC somata, recapitulating their organization *in vivo* between P5 and P7 **(Fig. 3C, 3I)** ^42,46,47^. In parallel, PF terminals emerged throughout the nascent ML, with increased VGLUT1/VGLUT2 colocalization **(Fig. 3C, 3J, 3L)**, consistent with the transient coexpression of both transporters by immature PFs as they establish contacts onto PC dendrites *in vivo* ^48^. Within the developing GCL, larger glutamatergic boutons expressing VGLUT1 and/or VGLUT2, consistent with maturing MF terminals ^49^, were also detected **(Fig. 3C)**. Inhibitory terminals contacted PC somata and developing dendrites, consistent with emerging MLI innervation **(Fig. 3D)**, while other VGAT+ terminals surrounded MF boutons within the GCL, indicating the concomitant assembly of cerebellar glomeruli containing GoC processes **(Fig. 3C, 3K)**.

At DIV21 (∼P14), PCs exhibited well-developed dendritic arbors **(Fig. 3E-3F)**. CF terminals were now predominantly localized along the primary dendrite, although some remained on the apical soma **(Fig. 3E, 3I)**, reflecting the ongoing somato-dendritic translocation ^50^. PF terminals became more abundant and were largely restricted to distal dendrites **(Fig. 3E, 3J)**, paralleling the marked expansion of PF innervation occurring *in vivo* during the corresponding developmental period ^51^. Inhibitory terminal density also increased throughout the somato-dendritic compartment **(Fig. 3F, K)**, consistent with extensive innervation of PCs by stellate and basket cells ^52^.

By DIV26, PCs displayed more elaborate dendritic arborization **(Fig. 3G-3H)**. CF terminals were largely absent from PC somata and concentrated along primary dendrites, as reflected by the significant increase in the dendritic-to-soma VGLUT2 density ratio **(Fig. 3I)**. PF and inhibitory inputs had reached distributions similar to those observed at DIV21 **(Fig. 3G-3H, 3J-3K)**, indicating stabilization of their mature spatial organization.

Together, these observations show that synaptic connectivity in hindbrain explants emerges through a coordinated developmental sequence closely resembling that observed *in vivo*. As PCs progressively mature, CFs transition from somatic innervation to their characteristic proximal dendritic territory, PFs emerge and expand over distal dendrites, and inhibitory and MF-associated circuits assemble in parallel. Thus, the mature organization of the explant is not simply an endpoint of prolonged culture but arises through the characteristic spatiotemporal progression of cerebellar microcircuit development.

### Hindbrain explants enable analysis of spontaneous olivo-cerebellar activity across development

Although the distribution of synaptic markers suggest that excitatory and inhibitory inputs mature appropriately, it does not prove whether cerebellar circuitry becomes functionally active. A defining feature of the mature olivo-cerebellar system is the generation of complex spikes (CSs) in PCs in response to the spontaneous activity of IONs. To investigate whether these characteristic physiological properties emerge in explants, we performed whole-cell patch-clamp recording of PCs and IONs from Igsf9-GFP explants at DIV12-13 (corresponding to ∼P5) and DIV28-30 (corresponding to ∼P21).

Current-clamp recordings in DIV30 explants revealed spontaneous spiking activity in both PCs and GFP+ IONs **(Fig. 4A–F)**. PCs exhibited characteristic CSs, consisting of a large sodium spike followed by spikelets superimposed to a depolarizing plateau, as described in acute slices and *in vivo* ^53^ **(Fig. 4A-4C)**. This indicates that IONs remain spontaneously active and are capable of eliciting the stereotypical all-or-none CF response in PCs. Consistent with this, IONs displayed spontaneous action potentials resembling those recorded in acute brainstem slices, characterized by a fast sodium spike followed by wavelets superimposed on an after-depolarization and a prolonged after-hyperpolarization phase ^12^ **(Fig. 4D-4E)**. Notably, the firing rate of IONs (0.08 Hz, IQR = 0.44 Hz) was comparable to values reported in brainstem acute slices from patch-clamp recordings ^54^, suggesting that ION remain spontaneously active in mature explants **(Fig. 4F)**.

**Figure 4:**
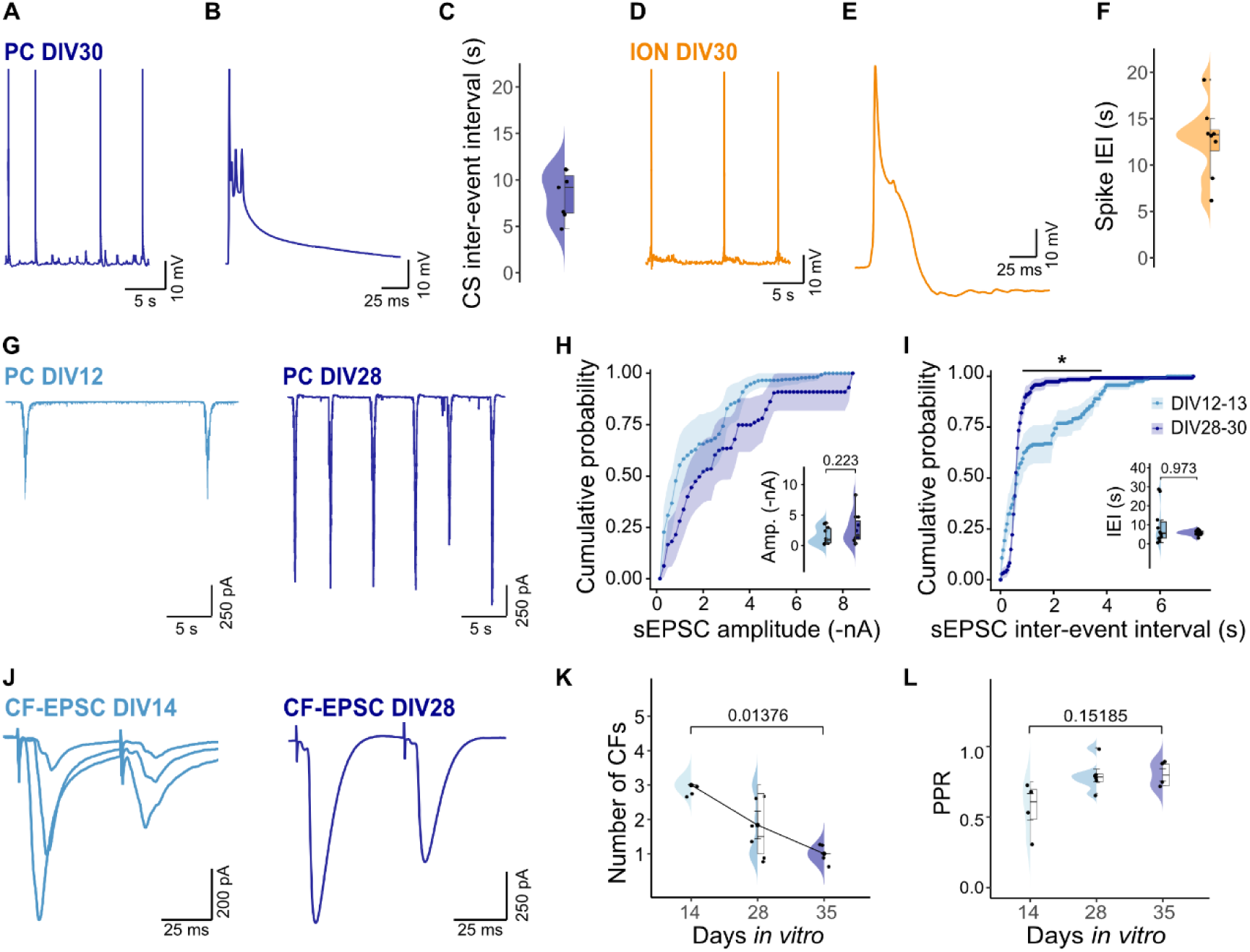
Hindbrain explants develop a spontaneously active olivo-cerebellar circuit. **(A)** Representative trace of current-clamp recording from DIV28 PC. **(B)** Expanded view of complex spike labelled (*) in (a). **(C)** Median IEI (ms) of complex spikes recorded from DIV28-30 PCs. Median (IQR) was 9170 ms (4025 ms); n=7. **(D)** Representative trace of current-clamp recording from DIV30 ION. Scale bar as in (A). **(E)** Expanded view of complex spike labelled (*) in (D) Scale bar as in (B). **(F)** IEI (ms) of complex spikes recorded from DIV30 IONs. Median (IQR) was 13260 ms (2267 ms); n = 8. **(G)** Representative traces of voltage-clamp gap-free recordings from PCs at DIV12-13 (light blue, left) and DIV28-30 (darkblue, right). **(H)** Cumulative distribution of large, CF-like sEPSCs events. Median (IQR) amplitudes were −953 pA (2377 pA) at DIV12–13 and −1631 pA (2988 pA) at DIV28–30. Violins correspond to the median large-events amplitude per cell. **(I)** Cumulative distribution of IEI (ms) of CF-like sEPSC. Median (IQR) IEI were 5599 ms (8454 ms) at DIV12–13 and 5819 ms (1636 ms) at DIV28–30. **(H, I)** DIV12-13 n = 10; DIV28-30 n = 11. Ribbon corresponds to SEM. Significance (*, p < 0.05) corresponds to pointwise Wilcoxon’s rank-sum test with Benjamini-Hochberg correction (FDR). **(J)** Representative voltage-clamp traces of evoked CF-EPSCs using paired-pulse stimulation at DIV14 (light blue, left) and DIV28 (dark blue, right). **(K)** Estimated number of CF inputs per cell over time. **(L)** Average CF-input PPR per cell over time. **(K, L)** Significance was determined using Kruskal-Wallis test (number of CF inputs p = 0.01726; PPRp = 0.1054) followed by Dunn’s pairwise comparisons with Bonferroni correction (values displayed on graph). DIV14 n = 4; DIV28 n = 6; DIV35 n = 4.

To assess the functional maturation of the olivo-cerebellar circuit, we performed voltage-clamp recordings from PCs at DIV12–13 and DIV28–30. Spontaneous events analysis revealed two amplitude-defined clusters **(Sup. Fig. 3A-2C)**: the nano- and pico-ampere range. The high-amplitude cluster, likely corresponds to spontaneous CF activity originating from IONs **(Fig. 4G)**, as these events were abolished by acute disconnection of the brainstem from the cerebellar plates **(Sup. Fig. 3D-2E)**. Between DIV12–13 and DIV28–30, they increased in both amplitude (∼70% increase) and frequency (∼60% decrease in inter-event interval) **(Fig. 4H-4I)**. At DIV28-30 their median inter-event interval was ∼5.8 s (IQR = 1.6 s), corresponding closely to frequencies reported *in vivo* ^55^. The smaller-amplitude events (-16.70 pA to -37.28 pA), likely corresponding to PF inputs and miniature activity from all glutamatergic afferents **(Sup. Fig. 3F-2H**).

Finally, we performed evoked recordings of CF-EPSCs in DIV14, DIV 28 and DIV35 explants using a paired-pulse stimulation protocol and identified CF inputs by their characteristic paired-pulse depression and their all-or-none nature **(Fig. 4J)**. In agreement with our previous findings ^32^, we found that the number of CF inputs per PC decreased over time **(Fig. 4K)**, reflecting postnatal CF refinement ^4^. Furthermore, the average paired-pulse ratio (PPR) reached larger values at DIV28-P35 **(Fig. 4L)** as reported *in vivo* ^56^.

Together, our results demonstrate that the olivo-cerebellar circuit matures and remains physiologically active in explants, proving the suitability of the system for the study of evoked but also spontaneous CF-PC signalling.

### Direct access to inferior olivary neurons enables morphological and long-range connectivity analysis in hindbrain explants

The preservation of spontaneous CF activity demonstrates that functional connectivity develops normally. Because both pre- and postsynaptic neurons remain directly accessible in explants, we next exploited this preparation to examine individual IONs and their long-range projections at single-cell resolution. We investigated whether the organization and morphology of IONs in Igsf9-GFP explants preserve typical *in vivo* features, including compact distribution of cell bodies around the midline, heterogeneous morphologies ranging from straight to highly tortuous “curly” forms, dendro-dendritic gap junction coupling, and contralateral projection of a single axon to the cerebellar plate and DCN ^57–62^.

At >DIV28, GFP+ IONs were bilaterally distributed along the midline (up to ∼230 µm laterally and, ∼650 µm in the antero-posterior axis) and largely exposed at the ventral surface, providing direct accessibility **(Fig. 5A)**. We implemented single-cell electroporation ^63^ to express tdTomato in individual IONs and identify their targets in the cerebellar plate. Each tdTomato+ ION extended a single axon that crossed the midline and projected contralaterally toward the cerebellar plate and DCN **(Fig. 5A, Sup. Fig. 4A).** Each tdTomato+ ION targeted 5 to 8 PCs (n = 5 IONs from 4 explants) **(Fig. 5B)**, consistent with previous *in vivo* single CF tracing studies and suggesting similar degree of circuit specificity and refinement ^64,65^.

**Figure 5:**
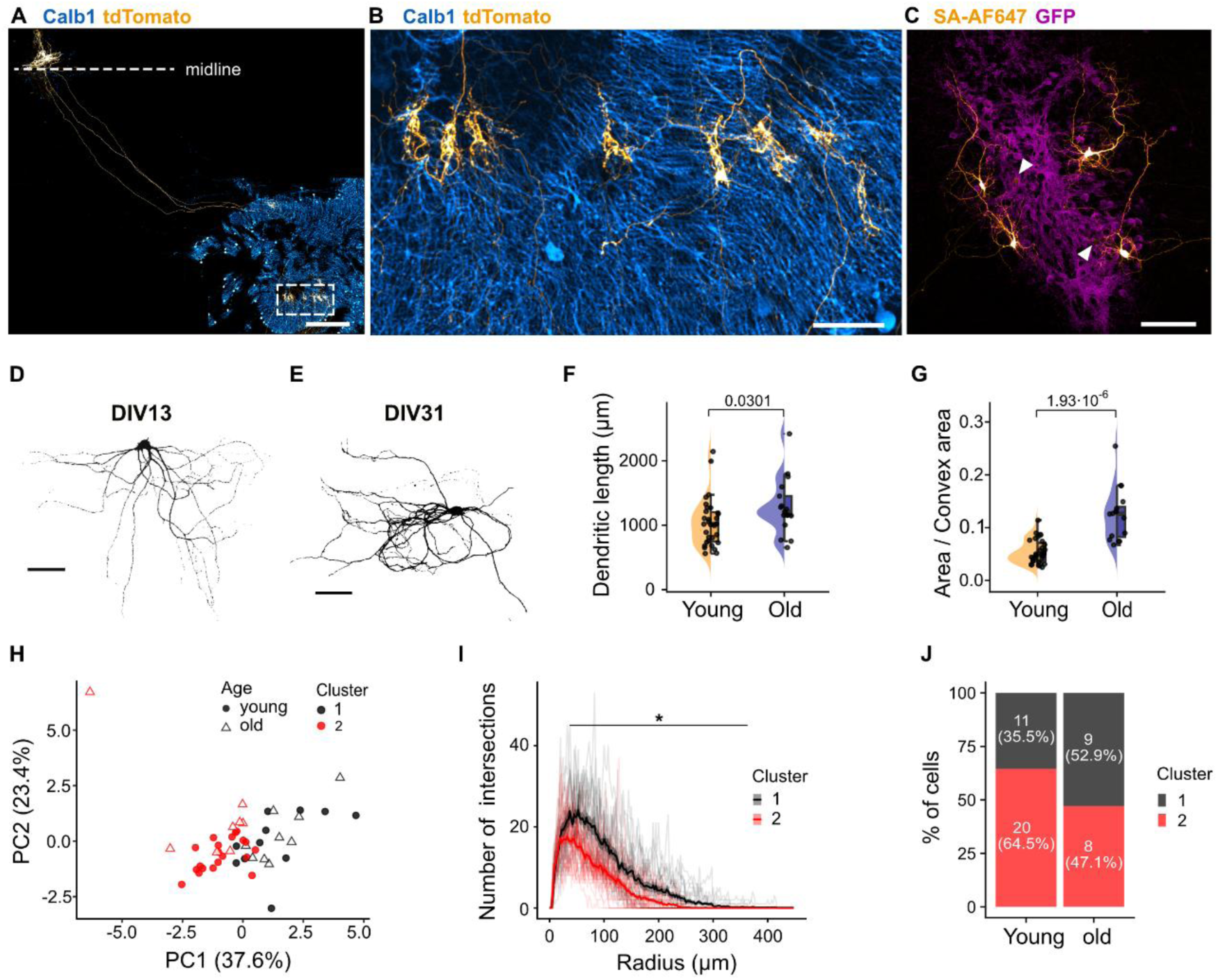
IONs undergo morphological maturation and establish characteristic connectivity with PCs in hindbrain explants. **(A)** Confocal mosaic image (MIP) showing two IONs at DIV30 labelled through SCE with tdTomato (yellow) with immunostaining for Calb1 (blue). CF cross the explant midline and innervate Calb1+ PC targets in the cerebellar plate. Scale bar = 300 µm. **(B)** Zoomed image of the PC targets in (A). Scale bar = 50 µm. **(C)** Confocal image (MIP) showing four biocytin-filled IONs at DIV13 (SA, orange) in the inferior olive region from an Igsf9-GFP explants (magenta). Scale bar = 100 µm. Arrowheads indicate the presence of putative IONs filled with biocytin through GAP junctions. **(D, E)** Thresholded images illustrating heterogeneous morphologies of IONs at DIV13 and DIV31. Scale bar = 50 µm. **(F)** Total dendritic length calculated from Sholl analysis at DIV12-13 (n = 31 cells from 12 explants, median (IQR) = 1015 µm (456 µm)), and DIV30 - 31 (n = 17 cells from 12 explants, median (IQR) = 1206 µm(304 µm)). **(G)** Dendritic field solidity (proportion of cell area covering the area of its convex polygon) at DIV12-13 (median (IQR) = 0.049 (0.034)) and DIV30-31 (median (IQR) = 0.126 (0.057)). **(F, G)** Displayed p-values correspond to Wilcoxon’s rank-sum test. **(H)** PCA of morphological features with cells projected into PCA space based on the first two principal components. Colors indicate k-means cluster assignment (k = 2), and shape indicates age group. Percentages on axes indicate the proportion of variance explained by each dimension. **(I)** Average Sholl profile per PCA cluster (mean trace, ribbon corresponds to SEM). Semi-transparent traces correspond to Sholl profiles of individual cells. Star region corresponds to the segment significantly different (p <0.05) between clusters. Significance was assessed by pointwise Wilcoxon’s rank-sum test with Benjamini - Hochberg correction (FDR) for each radius bin. **(J)** Proportion of cluster 1 and cluster 2 cells within each age group.

We next examined the morphology of IONs that were recorded through patch-clamp recordings at DIV12-13 or DIV28-DIV30, filled with biocytin and subsequently labelled with fluorescent streptavidin **(Fig. 5C)**. Heterogeneous morphologies were observed at both developmental stages. Moreover, a faint but highly specific fluorescent signal was observed from cells in the close proximity of the patched IONs **(Fig. 5C)**, suggesting that gap junction coupling is preserved in explants ^58^. Morphological analysis revealed a developmental increase in total dendritic length **(Fig. 5F)** and dendritic field solidity **(Fig. 5G)** indicating growth and increased complexity of ION dendrites over time.

To assess ION morphological heterogeneity in a more integrated and objective manner, we performed k-means clustering on principal component analysis (PCA) reduced data combining Sholl parameters (maximal dendritic length, total number of intersections normalized to the maximum length, Sholl radius with peak intersections, total dendritic length) with morphological descriptors (cell area, perimeter, solidity, circularity, roundness) **(Sup. Fig. 4B-F)**. Silhouette analysis identified k = 2 as the optimal clustering solution, revealing two ION morphological groups that were present across all ages **(Fig. 5H)** and differed in both cell size and shape **(Sup. Fig. 4C-D)**. Significant differences in average Sholl profiles between clusters **(Fig. 5I)** confirmed that the *in vivo* morphological diversity of IONs is preserved in explants. Although the distribution of cells between the two clusters was more balanced at DIV28-30, both curly and straight dendrites (along with intermediate shapes) were observed in both age groups **(Fig. 5J)**, indicating early postnatal diversification.

Together, these results show that IONs in explants retain the characteristic ION-PC connectivity stoichiometry while diversifying morphologically to generate dendritic configurations similar to those found *in vivo*.

### Gene expression regulation in PCs and IONs during circuit development

Having established that the explant system retains the morphological, functional and synaptic maturation of the olivo-cerebellar circuit, we next performed single-cell RNA sequencing of our multimodal PatchSeq data to determine whether these features are accompanied by the corresponding molecular programs ^66^ **(Fig. 6A, Sup. Fig. 4A)**.

**Figure 6:**
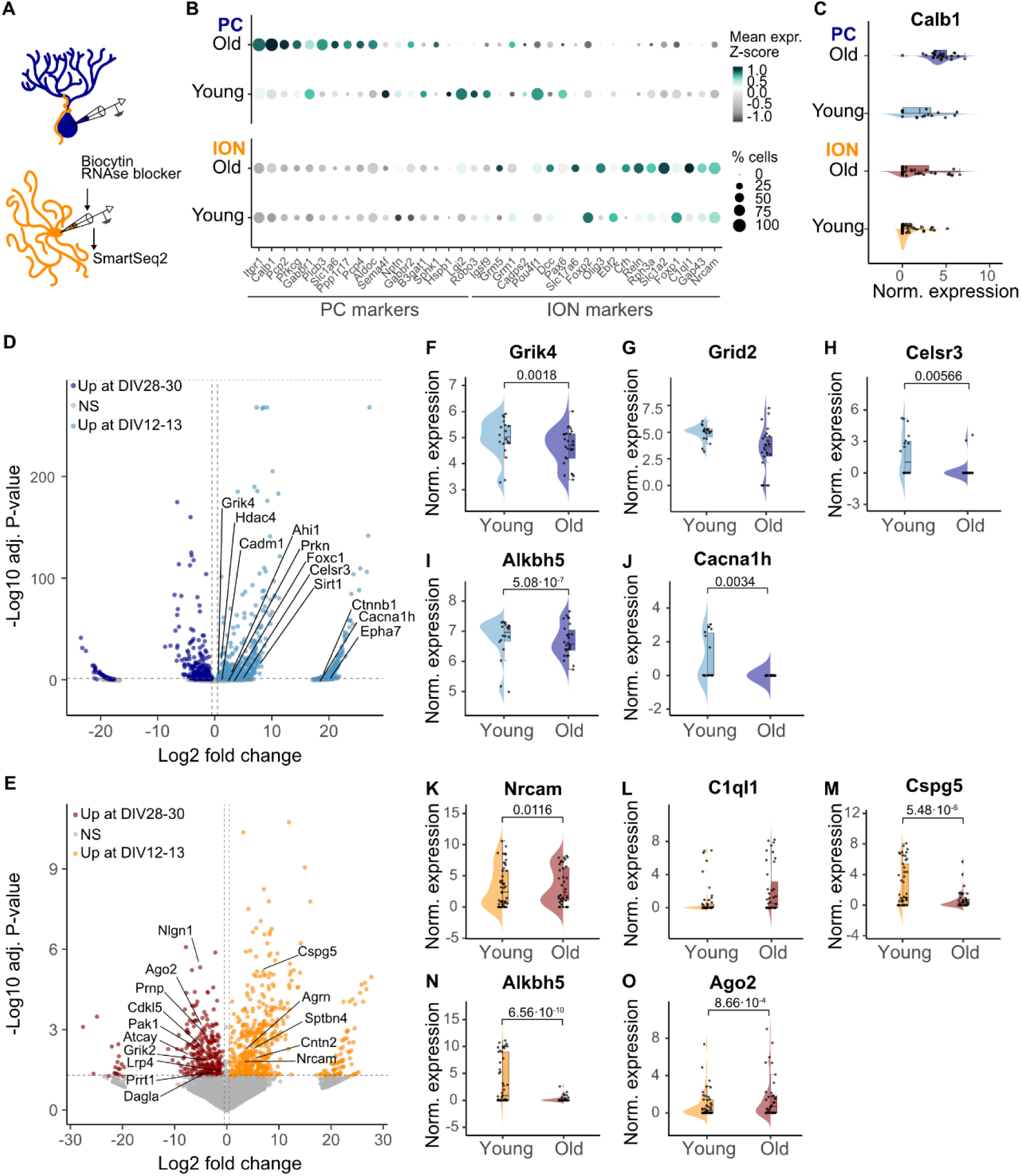
PCs and IONs retain their molecular identity and undergo transcriptional maturation in hindbrain explants. **(A)** Schematic representation of PatchSeq experimental setup in PCs and IONs. **(B)** Dotplot showing selected marker genes expression in PCs and IONs across development. Dot size corresponds to the percentage of expressing cells and color scale indicates the mean expression Z-score calculated per gene. Genes were ordered by relative enrichment in PCs vs. IONs. PC DIV12-13 n = 18 from 5 explants, DIV28-30 n = 26 from 12 explants; ION DIV12-13 n = 45 from 14 explants, DIV28-30 n = 39 from 18 explants. **(C)** Log-normalized expression of Calb1 in PCs and IONs at DIV12-13 and DIV28-30. **(D-E)** Volcano plot showing the differentially expressed genes (DESEq2’s LRT, adjusted p-value < 0.05) between DIV12-13 and DIV28-30 in PCs (D) and IONs (E). **(D)** Upregulated genes in DIV12-13 PCs: N = 1333 genes; downregulated genes in DIV12-13 PCs: N = 301 genes. **(E)** Upregulated genes in DIV12-13 IONs: N = 395 genes; downregulated genes in DIV12-13 IONs: N = 249 genes. **(F-O)** Log-normalized expression of differentially expressed GOIs in PCs (blue, F-J) and IONs (orange, K-O). Displayed values correspond to LRT adjusted p-values. **(F)** Grik4 log2-Fold Change (log2FC) = 0.918. **(G)** Grid2 log2FC = 0.232. **(H)** Celsr3 log2FC = 4.423. **(I)** Alkbh5 log2FC (in PCs) = 0.624. **(J)** Cacnca1h log2FC = 18.27. **(K)** Nrcam log2FC_= 3.22. **(L)** C1ql1 log2FC = -1.975. LRT adj. p-value = 0.119. **(M)** Cspg5 log2FC = 7.043. **(N)** Alkbh5 log2FC (in IONs) = 15.172. **(O)** Ago2 log2FC = -4.68.

We first validated the molecular identity of recorded neurons using a curated panel of PCs and IONs marker genes **(Fig. 6B, Sup. Fig. 5B-C)**. PC markers were selected based on previous transcriptomic characterization of the cerebellum ^67^ while ION markers were compiled from previous literature on olivo-cerebellar circuit development ^33,68–78^.

We next investigated developmental transcriptional changes between DIV12-13 and DIV28-DIV30. First, well-known genes such as *Calb1*, a shared marker for IONs and PCs, showed the expected developmental upregulation **(Fig. 6C)**. Known genes of interest (GOIs) for cerebellar development and olivo-cerebellar connectivity were found regulated during development ^52,69^. Furthermore, differential expression analysis (DESeq2’s likelyhood ratio test - LRT) followed by Gene ontology enrichment analysis ^79^ allowed the unbiased identification of up- or downregulated GOIs (adjusted LRT p < 0.05) associated with cerebellar function, synaptic maturation and neuronal activity **(Fig. 6D - 6E)**.

In PCs, our analysis revealed a developmental downregulation of genes associated with synaptic signaling, neuronal development or neuronal excitability, including the kainate receptor *Grik4*, the non-classical cadherin *Celsr3*, the RNA demethylase *Alkbh5* and the T-type Cav3.2 channel *Cacna1h* ^80–84^ **(Fig. 6F-J).** In contrast, *Grid2*, encoding the GluD2 receptor, remained stably expressed, consistent with its established ro le throughout PC maturation ^85^ **(Fig. 6G)**. Likewise, IONs exhibited developmental upregulation of genes involved in axon growth and synaptic development, including *Nrcam*, chondroitin sulfate proteoglycan (*Cspg5*), and *Alkbh5* **(Fig. 6K-N)**, while *C1ql1*, known to be directly involved in CF to PC synapse development, showed a trend towards higher expression at DIV28-30 ^69,81,86,87^ **(Fig. 6L).** The RISC component *Ago2* was found upregulated at DIV28-30 **(Fig. 6O)**, consistent with its role in PC morphogenesis ^88^. Beyond these established GOIs, we could identify the developmental regulation of genes related to cerebellar pathologies, such as *Ahi1*, *Ctnnb1*, *Sptbn4* or *Prkn* ^89–92^, highlighting the potential of the explant preparation for investigating disease associated molecular pathways.

Together, these results indicate that both PCs and IONs in explants retain molecular identity, while undergoing coordinated transcriptomic changes highly relevant for cell maturation and developmental circuit refinement.

### Microglia ramify and show interaction with PCs during development in explants

Glial cells, including microglia, play important role in neural circuit development ^36,93^. Thus, the usefulness of *in vitro* preparations critically depends on their ability to preserve glial cells characteristics, often heavily altered by slicing or dissociation procedures associated with culture models ^30^. We therefore examined whether microglia undergo the typical morphological transformation observed *in vivo* during development, in particular the increase in ramification and the decrease in soma size ^36^. To this aim, we immunolabelled microglia for Iba1 in fixed explants at DIV14, DIV17, DIV21 and DIV28 **(Fig. 7A)**. Sholl analysis revealed a progressive increase in microglia branching complexity, with both the peak number of intersections and total process length increasing over time **(Fig. 7B-7D)**. Similarly, branch number significantly increased throughout development **(Fig. 7E)**, indicating progressive ramification. Furthermore, soma size significantly decreased, consistent with a transition towards a less phagocytic, homeostatic phenotype **(Fig. 7F)**. Together, these data show that microglia in explants recapitulate a gradual morphological maturation transitioning from an amoeboid to a ramified state during development ^94,95^.

**Figure 7:**
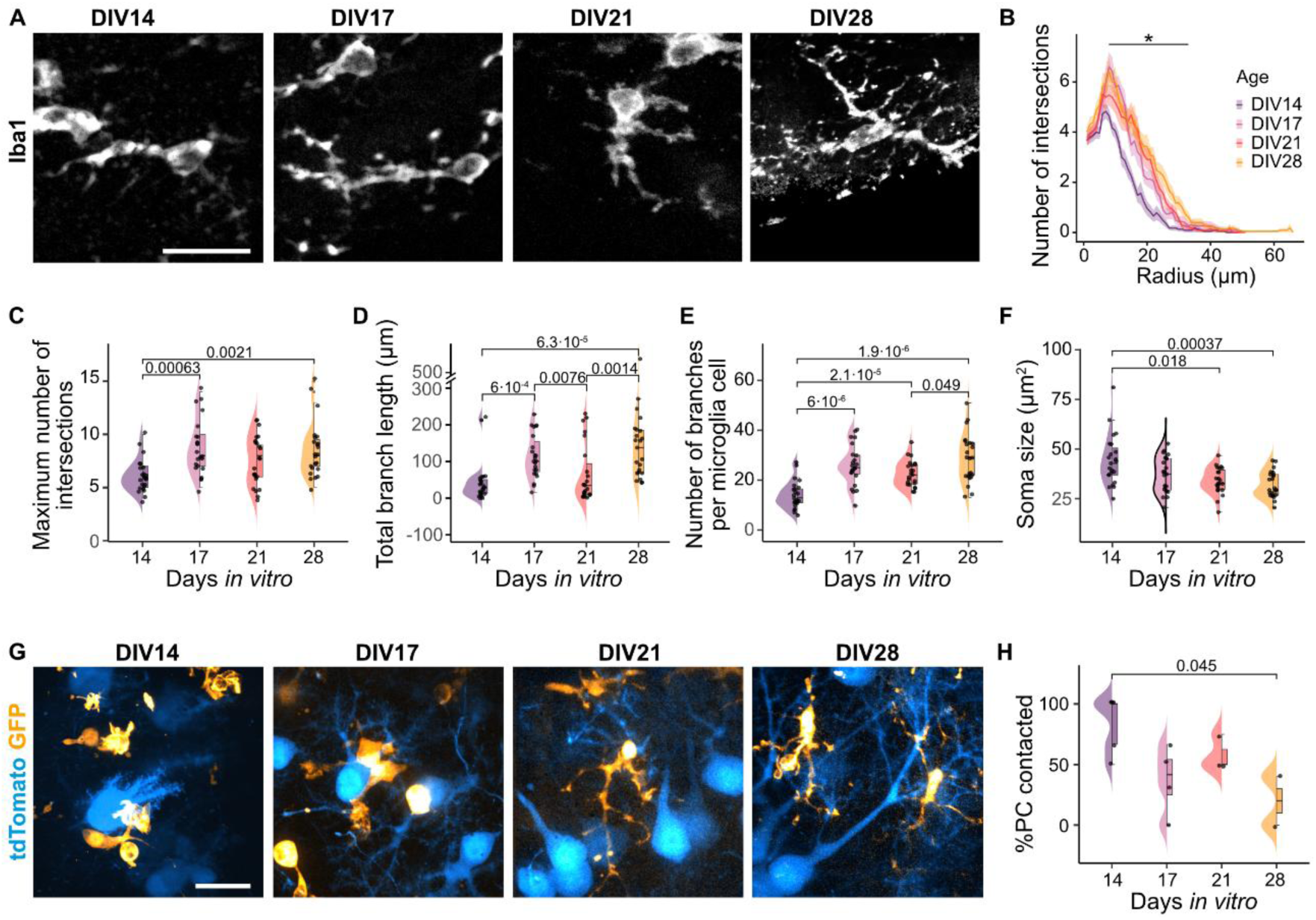
Microglia develop a ramified morphology and remain dynamically active in hindbrain explants. **(A)** Confocal images (MIP) of Iba1+ microglial cells at DIV14, DIV17, DIV21 and DIV28. Scale bars = 20 µm. **(B)** Average Sholl profile of DIV14 (purple), DIV17 (magenta), DIV21 (red) and DIV28 microglia (yellow). N = 23 cells per time point. Ribbon corresponds to SEM. Differences between age groups was assessed using Kruskal-Wallis test per radius with Benjamini-Hochberg p-value correction (FDR) followed by Dunn’s post-hoc comparisons and same p-value adjustment. Segment corresponds to the region significantly different between DIV14 and DIV28 (p < 0.05). Significant differences were also found for DIV14 versus DIV17 between 8 µm and 24 µm radius, DIV14 versus DIV21 between 13 µm and 28 µm and DIV17 versus DIV28 at 25 µm and 30 µm. **(C)** Maximum intersections found in a single radius in each age group. Kruskal-Wallis p = 0.000359. Median (IQR) was 6 (2) intersections at DIV14, 8 (3) at DIV17, 8 (3) at DIV21 and 8 (2.5) at DIV28. **(D)** Total branch length across development. Kruskal-Wallis p = 2.75·10^-6^. Median (IQR) was 29 µm (37 µm) at DIV14, 101 µm (82.5 µm) at DIV17, 35 µm (84 µm) at DIV21 and 138 µm (116 µm) at DIV28. **(E)** Number of branches per microglia cell across development. Welch-ANOVA p =1.91·10^-8^. Displayed values correspond to Games-Howell post-hoc test. Mean ± SEM was 14.1±1.16 branches at DIV14, 26±1.69 at DIV17, 22.6±1.019 at DIV21 and 28.8±2.03 at DIV28. **(F)** Microglia soma size per age. Kruskal-Wallis p = 6,23·10^-4^. Median (IQR) was 42.8 µm^2^ (13.57 µm^2^) at DIV14, 37.32 µm^2^ (15.94 µm^2^) at DIV17, 32.78 µm^2^ (9.66 µm^2^) at DIV21 and 29.36 µm (10.45 µm^2^) at DIV28. **(C, D, F)** Displayed values correspond to Dunn’s posthoc comparisons. **(G)** MIP of time-lapse imaging of microglia (Cx3CR1-GFP, green) and PC (tdTomato, red) imaged every 30 s for 10 min from DIV14 to DIV28. Top row (t _i_) corresponds to MIP of the confocal stack acquired at the first time point. Middle row (t_f_) corresponds to the MIP of the stack acquired in the last time point (t_21_). Bottom row (Area surveilled) corresponds to the MIP of all time points (t _1_ to t_21_). **(H)** Percentage of PCs in a field of view contacted by microglia. ANOVA p = 0.033. Displayed value corresponds to Tukey’s HSD test. Mean±SEM was 83.3±10.5 % at DIV14, 37.5±14.2% at DIV17, 58.3±8.33% at DIV21 and 20±20% at DIV28. **(I)** Total area explored by microglia during 10 min. Kruskal-Wallis test p = 0.209. Median (IQR) was 604 (311) µm^2^ at DIV14, 473 (61.6) µm^2^ at DIV17, 627 (167) µm^2^ at DIV21 and 404 (99.3) µm^2^ at DIV28. **(H, I)** DIV14 n = 6; DIV17 n = 5; DIV21 n = 4; DIV28 n = 5.

Microglia are inherently motile cells that continuously surveil their environment and interact with neurons. To monitor microglial dynamics and their interaction with PCs, we performed live imaging at DIV14, DIV17, DIV21 and DIV28 in Cx3CR1-GFP explants co-transduced with AAV9-CAG-flex-tdTomato and AAV9-CaMKIIα-Cre, which results in PCs sparse labeling. During the 10 minutes imaging sessions, microglial process movements could be clearly observed **(Video S1)**. Process movement could be observed at all ages **(Videos S2-4)**, while cell body movements was occasionally observed, predominately at early developmental stages. Qualitative observations of contact between microglia and PCs indicate that microglia processes activity follow the development of PC, contacting initially the soma and progressively the developing dendrites. Indeed, the proportion of PCs contacted by microglia at the somatic level decreased with age **(Fig. 7H)**.

Together, these observations show that hindbrain explants conserve microglial cells characteristics and offers accessibility to microglia dynamics alongside ongoing neuronal maturation, thus allowing detection of developmental changes in microglia-neuron interactions that are challenging to capture *in vivo*.

### Hindbrain explants permit longitudinal investigation of CF-PC synapse refinement at the cellular resolution

Finally, a noteworthy advantage of culture systems is the accessibility for longitudinal interrogation and manipulation of synaptic circuit development at the single-cell resolution. To illustrate the possibility to follow the morphological maturation of individual CFs or PCs in the explant system, individual PCs or IONs were labelled with tdTomato by single-cell electroporation. After 2-3 days, 3D stack images were acquired daily by spinning-disk microscopy. PCs exhibited a marked expansion of their dendritic arbor between DIV19 and DIV29 **(Fig. 8A)**. On the presynaptic side, a single ION axon branched into two CFs and contacted two putative PC targets between DIV20 and DIV22. From DIV25 onwards, one CF progressively retracted and disappeared by DIV28-29 **(Fig. 8B-8C)**, consistent with developmental CF competition and refinement ^64,96,97^.

**Figure 8:**
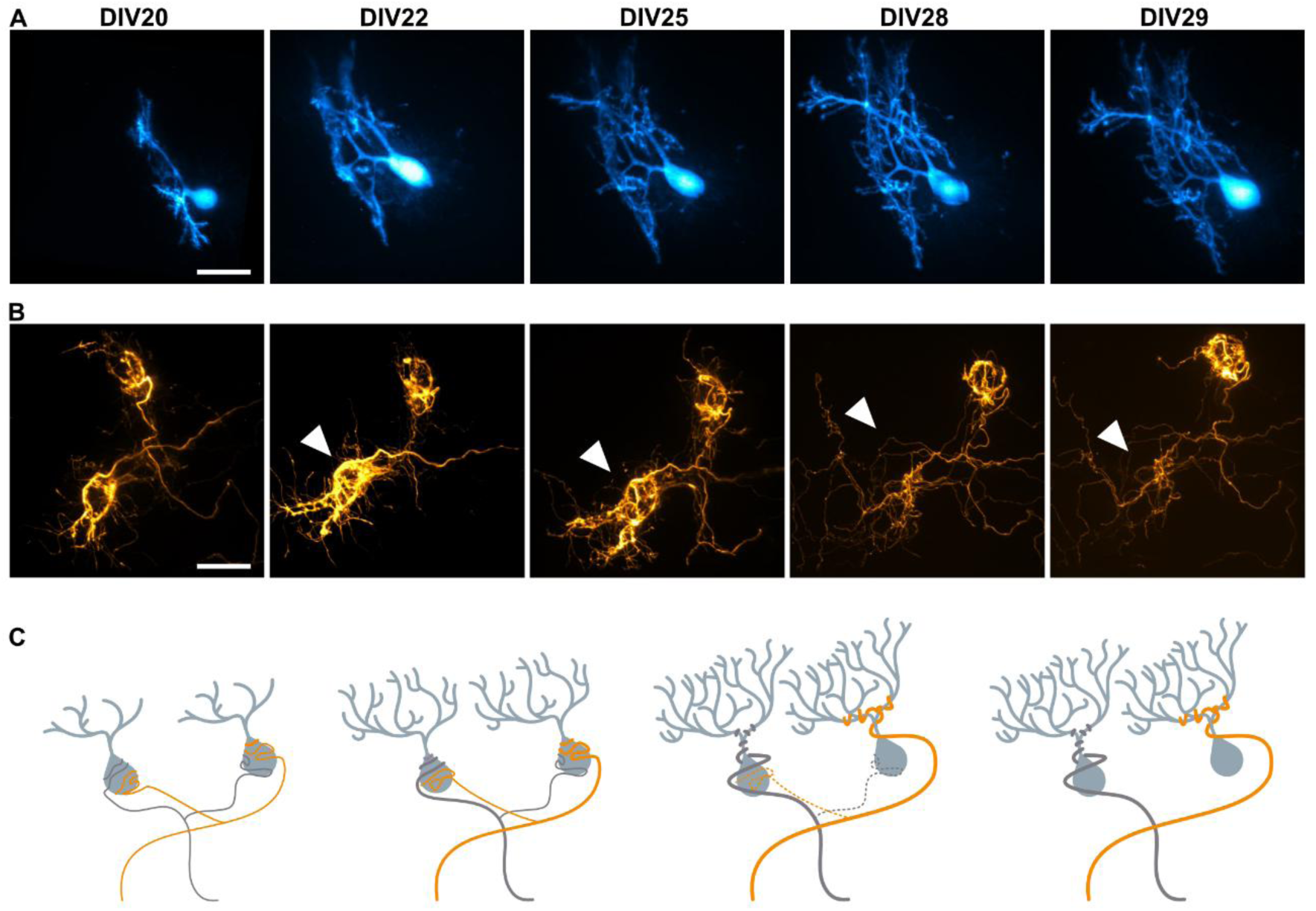
Hindbrain explants enable longitudinal analysis and manipulation of identified neurons during circuit development. (A) Morphological development of an individual PC in a hindbrain explant. (B) Strengthening of a CF branch (top) and simultaneouselimination froma different target (bottom, arrowhead) in a hindbrain explant. (A, B) Individual PC (A) or ION (B) were labelled through SCE using tdTomato and image from DIV20 (left) to DIV29 (right). Scale bar = 25 µm. (C) Schematic representation of two ION axons bifurcating and stabilizing different PC targets.

This observation demonstrates that hindbrain explants permit direct, longitudinal visualisation of CF branch remodelling and PC dendritic maturation at single-cell resolution. Compared to *in vivo* approaches, the explant preparation greatly simplifies high-resolution imaging and reconstruction of individual axons while capturing dynamic events that are otherwise difficult to follow over time ^97^.

## Discussion

Understanding how neural circuits assemble requires experimental models that preserve interactions between developing neuronal populations while providing access to the cellular and molecular events underlying synapse formation and refinement. Such preparations models are remarkably scarce for the mammalian CNS. Here, we establish hindbrain explants as a preparation that bridges this long-standing gap for the olivo-cerebellar circuit. We show that explants preserve the structural organization, spontaneous activity, developmental maturation and molecular identity of IONs and PCs while providing direct experimental access to both pre- and postsynaptic partners. By combining time-lapse imaging, targeted single-cell manipulation, electrophysiology and single-cell transcriptomics within the same intact preparation, we show that hindbrain explants constitute a versatile platform for investigating circuit assembly across molecular, cellular and functional scales.

One of the most striking observations is the remarkable fidelity with which olivo - cerebellar development proceeds despite prolonged culture conditions. Throughout several weeks *in vitro*, PCs and IONs undergo coordinated structural, physiological and transcriptional maturation that closely parallels *in vivo* development. CFs establish their characteristic pattern of innervation, undergo developmental refinement, and generate spontaneous CSs. Likewise, PCs molecularly diversify while maintaining their characteristic cytoarchitecture and synaptic organization. Collectively, these findings indicate that the core developmental program governing olivo-cerebellar assembly is remarkably robust and can unfold outside the intact brain, provided that reciprocal connectivity and spontaneous activity are maintained.

This finding has broader implications for understanding cerebellar development. Several studies have demonstrated that reciprocal interactions between pre- and postsynaptic partners are required for dendritic maturation, synapse elimination and circuit refinement in cerebellum and other brain areas ^31,32,98^. Our results indicate that preserving the complete olivo-cerebellar loop, including its endogenous patterns of spontaneous activity, is sufficient to support these developmental processes outside the intact brain. Rather than simply reproducing mature anatomy, hindbrain explants preserve a dynamically developing and physiologically active circuit, allowing developmental mechanisms to be studied as they unfold.

An important consequence of the accessibility of this model is the ability to integrate complementary approaches within the same preparation. We demonstrate targeted genetic manipulation of identified neurons, longitudinal imaging of individual CFs and PCs, and whole-cell recordings from both pre- and postsynaptic neurons. We further provide transcriptomic analysis revealing developmentally regulated genes, particularly for IONs, for which gene expression profiling was previously missing in the literature. Importantly, these approaches are not independent applications but complementary measurements that can be combined within the same experimental paradigm. This flexibility should facilitate mechanistic studies of activity-dependent circuit refinement and provide a framework for systematically linking gene expression, neuronal morphology, physiology and connectivity during development.

Beyond neurons, hindbrain explants also preserve a cellular environment compatible with the study of neuron-glia interactions. Organotypic slices are increasingly recognized to induce persistent inflammatory responses that influence synaptic remodeling and neuronal maturation ^30^. In contrast, microglia in hindbrain explants progressively acquire a ramified morphology while remaining dynamically motile, suggesting that the preparation maintains a relatively homeostatic environment throughout long-term culture. Although additional molecular characterization of glial states will be valuable, these observations indicate that hindbrain explants provide a unique opportunity to directly image and manipulate glial cells in homeostatic conditions to assess their contribution to cerebellar circuit assembly and plasticity.

Despite these strengths, hindbrain explants present certain limitations. The open-book configuration alters the three-dimensional geometry of the cerebellum, resulting in the lack of foliation, a patch-like PC distribution and dendritic arbors that extend within a broader two-dimensional plane than *in vivo*. Although spontaneous inferior olivary activity and PC CSs were readily detected, their firing frequency was lower than reported in anesthetized or awake mice ^12,99^, possibly resulting from the room-temperature recording conditions. In addition, the preparation lacks extracerebellar inputs and does not recapitulate systemic influences such as vascular, endocrine or behavioural modulation. Nevertheless, the remarkable preservation of structural, physiological and molecular developmental trajectories indicates that these factors are not essential for many of the core mechanisms governing olivo-cerebellar circuit assembly.

Collectively, our findings establish hindbrain explants as a physiologically faithful and experimentally accessible model of cerebellar development, whose potential is expanded by modern single-cell and multimodal technologies. By preserving an intact, spontaneously active, long-range mammalian circuit, this preparation enables the observation of identified neurons as they connect to their partners, manipulation of molecular pathways from early developmental stages, and correlation of changes in morphology and connectivity to neuronal physiology, transcriptional identity, and glial interactions. This model therefore bridges the gap between reductionist *in vitro* systems and technically demanding *in vivo* approaches. We anticipate that it will facilitate mechanistic studies of neural circuit assembly, neuron-glia interactions and cerebellar disease, while providing a scalable platform for genetic and pharmacological discovery.

## Materials and Methods

### Animals

All procedures involving mice and their care were conducted in accordance with the European guidelines for the care and use of laboratory animals, and the guidelines issued by the University of Bordeaux animal experimental committee (CE50; animal facilities authorizations #A33063940). Igsf9-eGFP mice (Tg(Igsf9-EGFP)JR10Gsat/Mmucd; REF PMID: 14586460) ^33^, kindly donated by Dr. Fekrije Selimi (Collège de France, Paris, France; MMRC n°030804-UCD) or Cx3CR1-GFP mice (JAX #005582) ^34^, were backcrossed onto the RjOrl:SWISS background (Janvier labs) to generate E14.5 embryos for olivo-cerebellar explants preparation.

### Hindbrain explants preparation and culture

Explants were prepared from E14.5 SWISS(x)Igsf-eGFP mice embryos, in which GFP is expressed under the ION-specific promoter *Igsf9*. Pregnant females were anesthetized through 5% isoflurane inhalation for 3 minutes and euthanised through cervical dislocation to quickly extract the uterus. Embryos were subsequently removed and placed in 4°C sterile dissection medium, containing (in mM): 25 glucose, 175 sucrose, 50 NaCl, 2.5 KCl, 0.5 CaCl_2_, 2 MgCl_2_, 0.28 MgSO_4_, 0.85 Na_2_HPO_4_, 2.7 NaHCO_3_, 0.66 KH_2_PO_4_ and 2 HEPES. Upon decapitation, brains were extracted and meninges were carefully removed **(Fig. 1A-B)**. The region between tectocerebellar and medullospinal junctions was isolated, and the two cerebellar plates were separated, obtaining an open-book configuration **(Fig. 1C)**. Explants were placed on Millicell culture inserts (Millipore PICM0RG50) and PTFE filter membranes (Millipore FHLC01300) with the dorsal side facing culture medium, containing: Basal Medium Eagle 50% (BME, Thermo 41010-026), Hank’s balanced salt solution 25% (HBSS, Gibco 14025-050), decomplemented horse serum 25% (Gibco 26050088), 25 mM glucose and 1 mM L-glutamine (Gibco 25030-024).

Explants were cultured at 35°C, 5% CO_2_ for up to 35 days in vitro (DIV) and medium was fully replaced every 2 days.

### Electrophysiological recordings and analysis

Whole-cell patch-clamp recordings were performed at room temperature (RT) using a Nikon Eclipse N1 upright microscope, equipped with infinity 3s camera (Teledyne Lumenera) driven by Metamorph® software (Molecular Devices), an apochromatic x60/1.0 NA water immersion objective (Nikon), and micromanipulators from Scientifica. Excitatory postsynaptic currents (EPSCs) and excitatory postsynaptic potentials (EPSPs) were recorded using a Multiclamp 700B amplifier (Axon Instruments, Molecular Devices), digitized at 20 kHz using a Digidata 1440A (Molecular Devices) and acquired using Clampex 10.7 software (Molecular Devices). Hindbrain explants were placed in the microscope chamber, continuously perfused with artificial cerebrospinal fluid (aCSF) bubbled with 95% O_2_/5% CO_2_ and containing (in mM): 25 glucose, 125 NaCl, 2.5 KCl, 2 CaCl_2_, 1 MgCl_2_, 1.25 Na_2_HPO_4_ and 25 NaHCO_3_. Micropipettes were obtained from 1 mm borosilicate capillaries (Harvard Apparatus) with a resistance of 5-7 MΩ using a vertical puller (Narishige). Internal solution for voltage-clamp recordings contained (in mM): 120 Cs-D-Gluconate, 10 HEPES, 10 BAPTA, 10 Na-P-Creatin, 3 TEA-Cl, 2 Na_2_ATP, 2 MgATP, 0.2 NaGTP and 13 biocytin (pH = 7,3; osmolarity adjusted to ∼300 mOsm). Internal solution for current-clamp recordings contained (in mM): 140 K-D-Gluconate, 6 KCl, 10 HEPES, 1 EGTA, 0.1 CaCl_2_, 5 MgCl_2_, 4 Na-ATP, 0.4 Na-GTP and 13 mM biocytin (pH = 7,3; osmolarity adjusted to ∼300 mOsm).

PCs were recorded from all cerebellar areas and were identified morphologically using DIC microscopy and fluorescence from Igsf9-GFP positive IONs CFs surrounding the target PCs. To isolate evoked CF events in PC (eCF-EPSC), 20µM bicuculline was added to the aCSF to block inhibitory synaptic transmission. A bipolar electrode placed in a borosilicate theta glass capillary filled with aCSF was placed in the GCL. Paired-pulse stimulation was applied to differentiate depressed CF-EPSC and facilitated PF-EPSC ^100^ (pulse duration range = 0.2 – 0.5 ms; inter-stimulus interval = 50 ms) using an ISO-Stim-01D isolated stimulator (NPI Electronic). To ensure the recruitment of all CFs connected to the recorded PC, the bipolar electrode position was adjusted and paired pulses were applied with increasing intensity in each location (from 10V to 95V). PC membrane potential was clamped at -70 mV for DIV12-13 explants, -20 mV for DIV28 and -10 mV for DIV30-35 for recording CF-EPSCs.

All analyses of evoked CF-EPSCs were conducted in a blind manner using Clampfit for event detection and a custom R script for subsequent quantification. To estimate the number of CFs connected to the recorded PC, ∼100 sweeps from pair-pulse stimulation were concatenated; the amplitudes of the first and second peaks (P1 and P2) relatively to the baseline were quantified in Clampfit (Molecular Devices, version 11.4.3). PPR (P2/P1) was calculated to discriminate CF-EPSCs from PF-EPSCs. Kernel Density Estimation (“density” function, R Core Team, 2013) with a Gaussian kernel function was employed to study the distribution of CF P1 amplitudes. Bandwidth was determined using pilot estimation of derivatives ^101^. P1 density function was thresholded using 1/5 of the standard deviation as a rule to discard outlier sweeps. The range of a given CF input was defined by calculating the local minima of the density function according to the second derivative rule. The range of P1 amplitudes belonging to each interval between local minima were averaged, as well as their paired P2.

To record spontaneous spikes in IONs or PCs, continuous (gap-free) 2 minutes recordings were performed in current-clamp (10 kHz Bessel filter) and voltage clamp mode (2 kHz Bessel filter, -70 mV holding potential). In current clamp recordings, or IONs spikes were detected and analysed using Clampfit 11.4.3 (Molecular Devices) threshold search for positive-going events (threshold at -10 mV). CSs in PCs were identified based on their typical shape consisting of a large sodium spike followed by spikelets superimposed to a long-lasting depolarizing plateau. Typical ION spikes were characterized by a fast sodium spike followed by wavelets superimposed on an after-depolarization and a prolonged after-hyperpolarization phase. The median inter-event interval (IEI) from all the events in a total of 4 minutes recording was calculated per cell. In voltage-clamp recordings, baseline was manually adjusted to 0 pA. Spontaneous EPSCs were detected using threshold search for negative-going events, adjusting the threshold to 5 times the standard deviation of the baseline (1 ms noise rejection, 1ms pre-trigger and post-trigger length). For each cell, the distribution of amplitudes of all the detected events was examined, identifying two clusters: large, CF-like currents, and small events. Each cluster of events was then analysed separately. The empirical cumulative distribution function (ECDF) was calculated for amplitude and IEI per cell. Bins were defined according to values range and number of events for each condition.

### PatchSeq sample processing and data analysis

#### Sample acquisition and sequencing

For Patch-seq sample acquisition, 0.5% RiboLock RNase Inhibitor (40 U µL⁻¹; Thermo Fisher Scientific) was added to the internal solution, and the osmolarity of the aCSF was adjusted to approximately 345 mmol kg⁻¹ using D-sorbitol. Upon completion of the electrophysiological recording protocol, the recording electrode was slowly withdrawn to obtain an outside-out configuration. The cytoplasmic content collected in the recording pipette was transferred into a PCR tube for cDNA synthesis according to the Smart-seq2 protocol ^66^.

cDNA libraries were pre-amplified by PCR and purified using AMPure XP magnetic beads. Sequencing libraries were subsequently prepared using the Nextera XT DNA Library Preparation Kit (Illumina). cDNA was subjected to transposase-mediated tagmentation, which simultaneously fragmented the cDNA and introduced adapter sequences. Following tagmentation, sequencing adapters and index sequences required for cluster generation and sample identification were added, and the libraries were amplified by PCR. After purification, library quality and fragment-size distribution were assessed by capillary electrophoresis using a LabChip GX Touch HT system (PerkinElmer), and library concentrations were quantified by qPCR. A unique index was incorporated into each sample during library preparation to enable multiplexing. Libraries were selected for sequencing based on their concentration and fragment-size distribution, pooled, and sequenced on a NextSeq 2000 platform (Illumina), targeting approximately 10 million reads per cell.

### PatchSeq analysis

#### Reads alignment

Adapter sequences were removed from raw reads using Cutadapt 1.18 ^102^ and fastp 0.20.0 ^103^. Quality of the pre-processed reads before mapping was assessed with FastQC version 0.11.5 ^104^. The splice aware aligner STAR 2.7.10a was used to map reads to the reference genome GRCm39 and to perform read counting at the gene level using default settings ^105^. Multi-mapping reads were discarded.

#### Quality controls

The total number of counts and total number of detected genes were evaluated, and outlier cells were discarded. Both parameters were also evaluated after subsetting datasets according to known experimental conditions to evaluate possible batch effects. Low-quality cells (less than 10000 total counts or more than 10000 detected genes) were excluded from downstream analyses. Counts were normalized using logarithmic normalization from Seurat (v5) ^106,107^. The expression of large, curated panels of PC and ION marker genes were evaluated and cells with no detected counts for any of the markers were discarded.

#### Differential expression and pathway analysis

Differential expression was assessed using Likelihood Ratio Test (LRT) in DESeq2 v1.42.1 ^108^ using explant age as the main test parameter. Pathway analysis of up- and downregulated genes was performed according to Biological Process, Cell Component and Molecular Function Gene Ontology (GO) terms enrichment using EnrichR ^79^. Genes of interest (GOI) were extracted from the enriched GO-terms and the panel was curated using the relevant bibliography.

### Single-cell electroporation

Explants were placed in the microscope chamber filled with sterile aCSF, containing (in mM): 130 NaCl, 2.5 KCl, 2.2 CaCl_2_, 1.5 MgCl_2_, 10 HEPES, 10 glucose. Micropipettes were filled with SCE solution, containing 9,5% TE buffer and 6.5 ng·µl^-1^ of pCAG-tdTomato plasmid in K-D-Gluconate based intracellular solution. IONs were identified by Igsf9-GFP positive fluorescence. Electroporation was performed after approaching the targeted cell and applying 4 consecutive pulses of -2,5 V, 25 ms duration, at 1 Hz. Explants were incubated for minimum 48h before imaging or fixation to allow constructs expression.

### Immunostaining, imaging and image analysis

### Immunostaining

Full, unsliced explants were used for all imaged shown except microglia images in Fig. 7A, for which transversal cryosections were acquired. For immunostaining, explants were fixed in 4% paraformaldehyde – 4% sucrose at 4°C overnight and free aldehyde groups were then quenched with NH_4_Cl 50 mM in PBS during 30 minutes at RT. Tissue was permeabilized using Triton X-100 0.5% in PBS for 15 minutes at RT. Non-specific binding was blocked by incubating samples in bovine serum albumin (BSA) 1% - 0.5% Triton X-100 in PBS during 1.5 h at RT. Exceptionally, blocking was performed during 24 h at 4°C for Aldolase C immunostaining. Proteins of interest were labelled using specific primary antibodies in BSA 1%-PBS at 4°C overnight, followed by fluorophores-conjugated secondary antibodies at 1 mg·ml^-1^ during 24 h at 4°C and 1 h at RT. Biocytin was revealed using streptavidin (SA) conjugated to AF488 or AF647 1 mg·ml^-1^. Samples were mounted on glass slides using Fluoromount with or without DAPI (Merck).

### Fixed samples imaging

Macroscopic images of embryonic brains were obtained using an Olympus SZX10 stereomicroscope equipped with an Olympus DP23 camera and cellSensEntry software. Confocal mosaics were obtained using a Zeiss Cell Discoverer 7 microscope with a 20X / 0.7 NA air objective and GaASP-PMT detector. Images of PC synaptic markers were acquired on a Leica DM6 CFS TCS SP8 confocal microscope using a 63X / 1.4 NA oil objective at 400 Hz scanning frequency and a pinhole opened to 1 time the Airy disk. Pixel size was set at 70-80 nm and z-step size was 0.3 µm. Confocal stacks of biocytin-filled IONs were obtained using the same system with a 40X / 1.3 NA oil objective. Pixel size was 160-162 nm and z-step size was 0.5 μm. Images of microglia were acquired using aa Nikon Eclipse FN-1 upright microscope, equipped with a Yokogawa CSU-22 spinning disk confocal scan head. Excitation was delivered via a laser bench (Errol) equipped with 491 nm, 561 nm, and 647 nm solid-state lasers. Emitted fluorescence was filtered through a multi-band emission filter wheel (447/460, 525/550, 535/545, 600/650 nm) controlled by a Lambda 10-3 controller (Sutter Instrument) and detected using an EMCCD camera (Rolera EM-C2, QImaging) with a 1004 × 1002 pixels array and 6.5 μm pixel pitch. Image acquisition and hardware synchronisation were managed using Metamorph Infinity software (v7.10, Molecular Devices). Z-stacks were acquired using a piezoelectric scanner (P-725.1CDE2, Physik Instrumente) and spanned 50 μm in depth with 0.3 μm axial intervals using a Nikon 60X/1.4 NA oil immersion objective.

### Live imaging acquisition

To enable repeated imaging of the same sample, sessions were limited to 30 minutes to reduce phototoxicity and contamination. Explants were placed in a sterile imaging chamber containing pre-warmed, sterile aCSF. Z-stacks spanning 40 μm were acquired at 1 μm z-step intervals every 30 seconds for 10 minutes using a Nikon Apo 60×/1.0 NA water dipping objective in the spinning disk system described above.

### Image analysis

All image analysis was performed in FIJI ^109^. To quantify PC afferent markers during development, the number of VGLUT2, VGLUT1, VGAT and colocalized puncta per µm^2^ was measured using the ImageJ plugin Puncta Analyzer by B. Wark (https://github.com/toddstavish/puncta-analyzer)^110^. Briefly, maximum intensity projection (MIP) images were obtained from 2 z-planes containing soma and dendrites. A median blur filter with 1-pixel radius was applied to all images to improve puncta detection and remove noise. For soma, regions of interest (ROI) were manually drawn around each soma based on the Calb1 immunolabelling. For dendrites, 3 representative ROIs of 20x20 µm were selected in the dendritic tree region based on Calb1 immunolabelling. Of note, the absence of dendritic tree in DIV7 explants did not allow the measurement of dendritic puncta at this stage.

To analyze ION morphology, a MIP was generated from each z-stack and biocytin signal was thesholded to obtain a binary image. A single ROI defining the cell contour was defined using *Analyze particles* and merging the resulting ROIs. Shape descriptors were then calculated from this ROI (cell area, perimeter, solidity, circularity and roundness). Sholl analysis was performed using SNT plugin on the thresholded image ^111,112^, setting manually a ROI to define the cell soma as a start point, and using 2.5 μm step size and automatic primary branches detection. Non-redundant variables from shape descriptors and sholl analysis were used to compute a PCA (maximal dendritic length, total intersections normalized to the maximum length, Sholl radius with peak intersections and total dendritic length, cell area, perimeter, solidity, circularity and roundness; all of them with a correlation between variables of < 40%). Cells were clustered using a k-means approach with *k* = 2. Optimal k value was defined attending to the average silhouette width obtained for each value of k from 1 to 10.

To analyze microglia morphology from fixed samples, a MIP was generated from each Z-stack. Each soma was manually traced three times to calculate an average area and corresponding effective radius. Microglia processes were traced with the Simple Neurite Tracer (SNT), and Sholl analysis was performed using 1 μm intervals between circles centred at the root centroid. To exclude the soma area from length estimates, the product of radius and the number of first-bin intersections was subtracted from the total processes’ length.

To quantify physical contacts between microglia and PCs, a MIP was created for each timepoint from each 4D dataset (x-y-z-t). The resulting 2D time series was temporally projected to identify candidate contacts, defined as spatiotemporally localised overlaps between GFP+ microglia and tdTomato+ PCs. Contacts were validated only if overlap occurred within the same z-plane with clear morphological continuity in the 3D z-stack. To quantify the dynamic surveillance area of microglial processes, we analysed 10-minute time-lapse recordings of GFP+ microglia. For each recording, a MIP was generated to obtain a 2D image for each timepoint. These were subsequently temporally projected to generate a z-t-stack single image capturing the full extent of process movement over the 10-minute window. The first frame was subtracted from the z-t-stack image. The resulting differential image was thresholded to quantify the area surveiled.

**Table 1:**
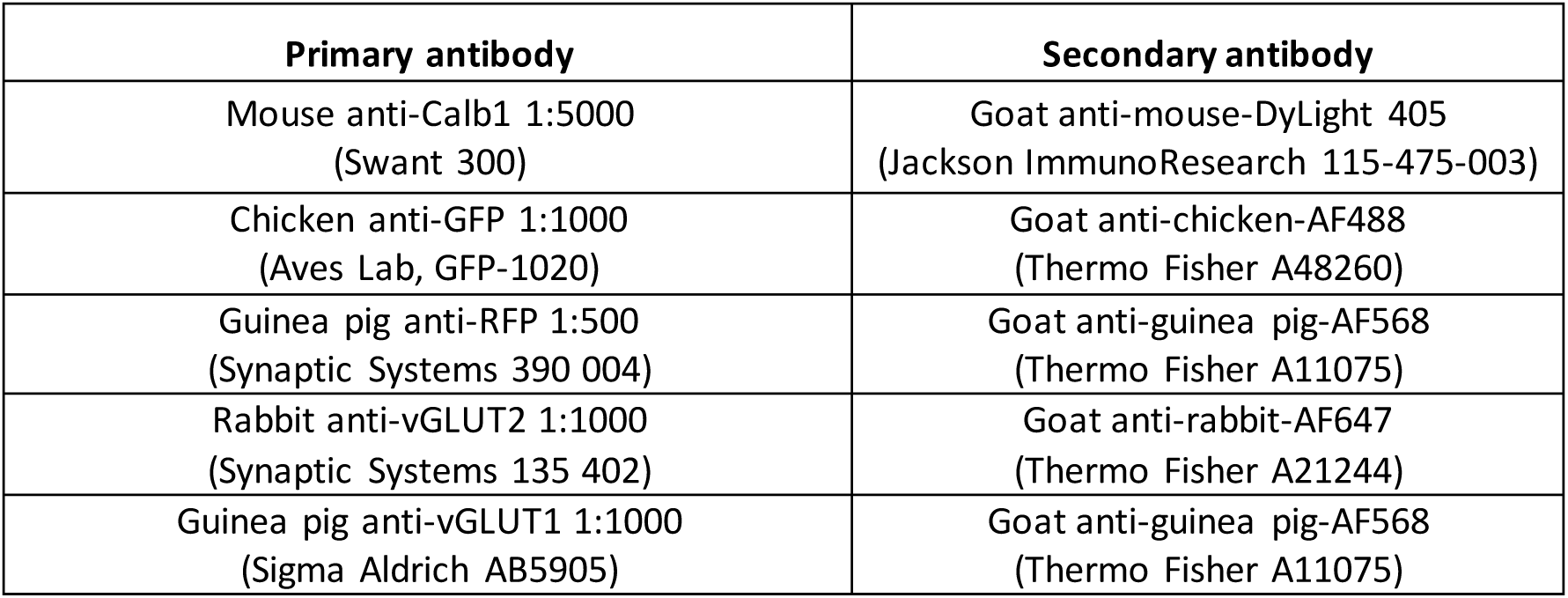
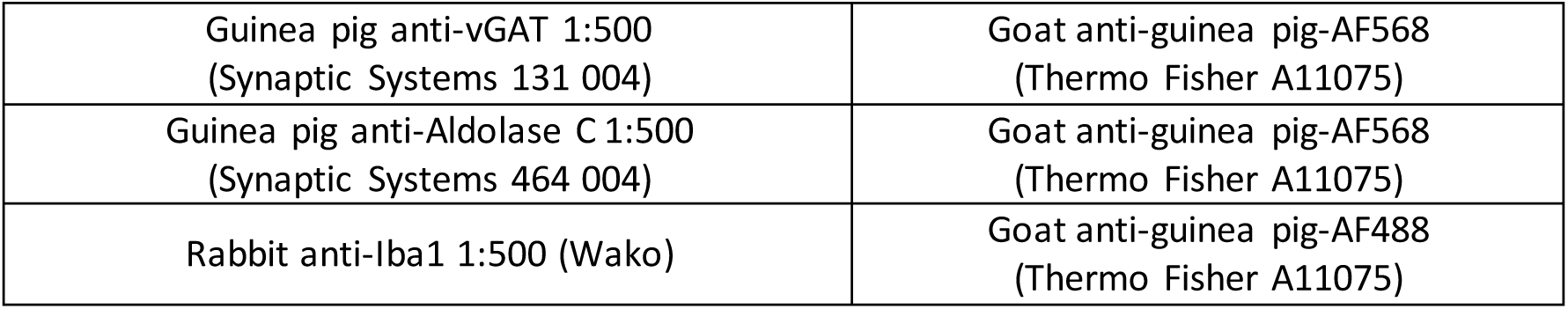
Primary and secondary antibodies combinations used.

### Transmission electron microscopy (TEM)

DIV30 explants (n = 2 explants) were fixed overnight at 4°C in 4% PFA and 1% glutaraldehyde in PBS. Explants were microdissected to retain only the cereb ellar plates. Samples were then incubated for 1 hour on ice in 1.5% OsO₄ and 1.5% potassium ferrocyanide (n° 20150 and 19150, respectively; Electron Microscopy Sciences, EMS, Hatfield, PA, USA) diluted in deionized water. Samples were dehydrated through graded ethanol solutions (70%, 90%, 95%, and 100%; 10 minutes each), followed by a 10 minutes incubation in 100% acetone. Tissues were then incubated for 1 h in a 1:1 mixture of acetone and Epon resin (Embed-812, EMS), followed by incubation in 100% Epon resin for 12 hours and subsequent polymerization at 60°C for 48 hours. Trapezoidal blocks containing cerebellar plate were trimmed in the transverse plane. Ultra-thin sections (70 nm) were obtained using an ultramicrotome (EM UC7, Leica, Germany) with a 35° diamond knife (Diatome, Switzerland) and collected on bare 150-mesh copper grids (G150-Cu, Delta Microscopy). TEM imaging was performed in high-contrast mode using a H7650 transmission electron microscope (Hitachi) operated at 80 kV and equipped with an Orius SC1000 CCD camera (11MPx, GATAN, Ametek, United-States). Acquisitions were performed by selecting regions located near PC as well as within the granule cell layer. Synapses were randomly imaged at magnifications ranging from 10,000× to 70,000× by acquiring every synaptic profile encountered until at least 30 synapses displaying a complete contour and clearly distinguishable ultrastructural features.

### Statistic comparisons and graphs

All statistical comparisons and graphs were performed in R using the packages *rstatistix* and *ggplot2*, respectively. Outliers were filtered using the 1.5xIQR Tukey criterion. For two-groups comparisons, normality and homoscedasticity were assessed using Shapiro-Wilk and Levene tests, respectively. t-test was used for normal data, and Welch correction was applied for non-homoscedastic sets. For non-normal data, Wilcoxon test was used.

For more than two groups comparisons, normality was examined with a Shapiro-Wilk test on the residuals of the ANOVA model. Homoscedasticity was checked with Levene test on the data. ANOVA followed by Tukey’s HSD pairwise comparisons was used for normal, homoscedastic data. Welch-ANOVA was used for normal, non-homoscedastic data, followed by Games-Howell post-hoc comparisons. Non-normal data was tested using Kruskal-Wallis test followed by Dunn’s pairwise comparisons.

## Resource availability

### Lead contact

Further information and requests for resources should be directed to and will be fulfilled by the lead contact, Mathieu Letellier.

### Materials availability

This study did not generate new unique reagents.

### Data and code availability

Patch-seq data generated in this study have been deposited in the ArrayExpress collection of EMBL-EBI under accession number E-MTAB-17525. The dataset is currently under embargo and will be made publicly available upon publication of the peer-reviewed article.

The custom R script used to identify and quantify CF inputs based on EPSC amplitude analysis is available at https://zenodo.org/records/22092907.

The custom R script used to perform differential gene expression analyses from PatchSeq datasets is available at https://zenodo.org/records/22092942.

Any additional information required to reanalyze the data reported in this paper is available from the lead contact upon request.

## Acknowledgments

This work received funding from the Centre National de la Recherche Scientifique, Agence Nationale pour la Recherche (ANR-22-CE16-0030-01), Fondation pour la Recherche Médicale (ECO202106013762 and FDT202404018274), Fondation pour la Recherche sur le Cerveau (FRC 2022-262421) and the University of Bordeaux (GPR BRAIN_2030). We thank the animal facility of the Bordeaux University (P. Costet, C. Martin, H. El-Houssini, C. Louis, E. Morillon, M. Fevre). We thank F. Cordelières, M. Fernandez-Monreal and S. Marais at the Bordeaux Imaging Center a service unit of the CNRS-INSERM and Bordeaux University, member of the national infrastructure France BioImaging supported by the French National Research Agency (ANR-24-INBS-0005 FBI BIOGEN). We also thank V. Demais (Strasbourg University) for helping with the interpretation of TEM images.

## Authors contributions

Conceptualization, E.B-B and M.L; formal analysis, E.B-B, C.T. and M.S-M; methodology, E.B-B, C.D., A.F., E.A. and M.L.; software, E.B-B and A.F.; investigation, E.B-B, C.T., M.S-M, C.D, L.F., T.V. and M.L.; data curation, E.B-B and M.L.; visualization, E.B-B and M.L.; writing – original draft, E.B-B and M.L.; writing – review & editing, E.B-B, C.T., M.S-M, C.D., L.F., T.V., A.F., E.A. and M.L.; funding acquisition, E.B-B, E.A. and M.L.; project administration, M.L.; resources, A.F. and M.L.; and supervision, E.A. and M.L.

## Supplementary figures

**Supplementary Figure 1:**
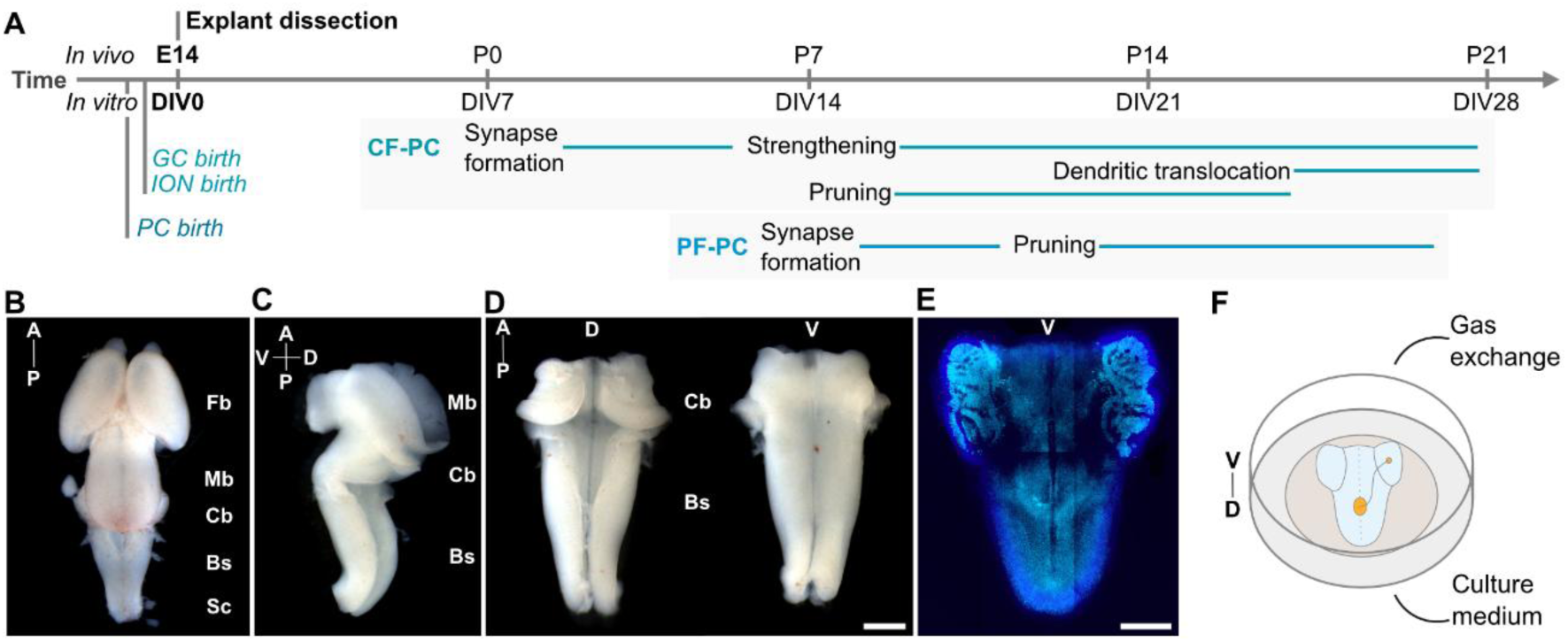
Hindbrain explants preparation and culture. **(A)** Developmental timeline recapitulating key embryonic (E) and postnatal events (P) in the cerebellar cortex, with equivalent developmental stages in explants (days in vitro, DIV). Explants are prepared from E14.5 mouse embryos, posterior to PC, ION and GC birth and can be maintained for several weeks covering the entire period of olivo-cerebellar circuit assembly. **(B)** Dorsal view of E14.5 mouse brain. **(C)** Lateral view of midbrain, cerebellum and brainstem at E14.5 mouse without meninges. **(D)** Dorsal (left) and ventral (right) views of an isolated hindbraincomprising cerebellumand brainstemprior culturing. Scale bar = 500 µm. **(E)** Epifluorescence image of a fixed hindbrain explant immunolabelled for Calb1 (cyan) and stained with DAPI (blue). Scale bar = 1 mm. **(B, C, D, E)** Fb: forebrain, Mb: midbrain, Cb: cerebellum, Bs: brainstem, Sc: spinal cord, A: anterior, P: Posterior, V: ventral, D: dorsal. **(F)** Schematic representation of an explant plated in a culture insert (ventral up).

**Supplementary Figure 2:**
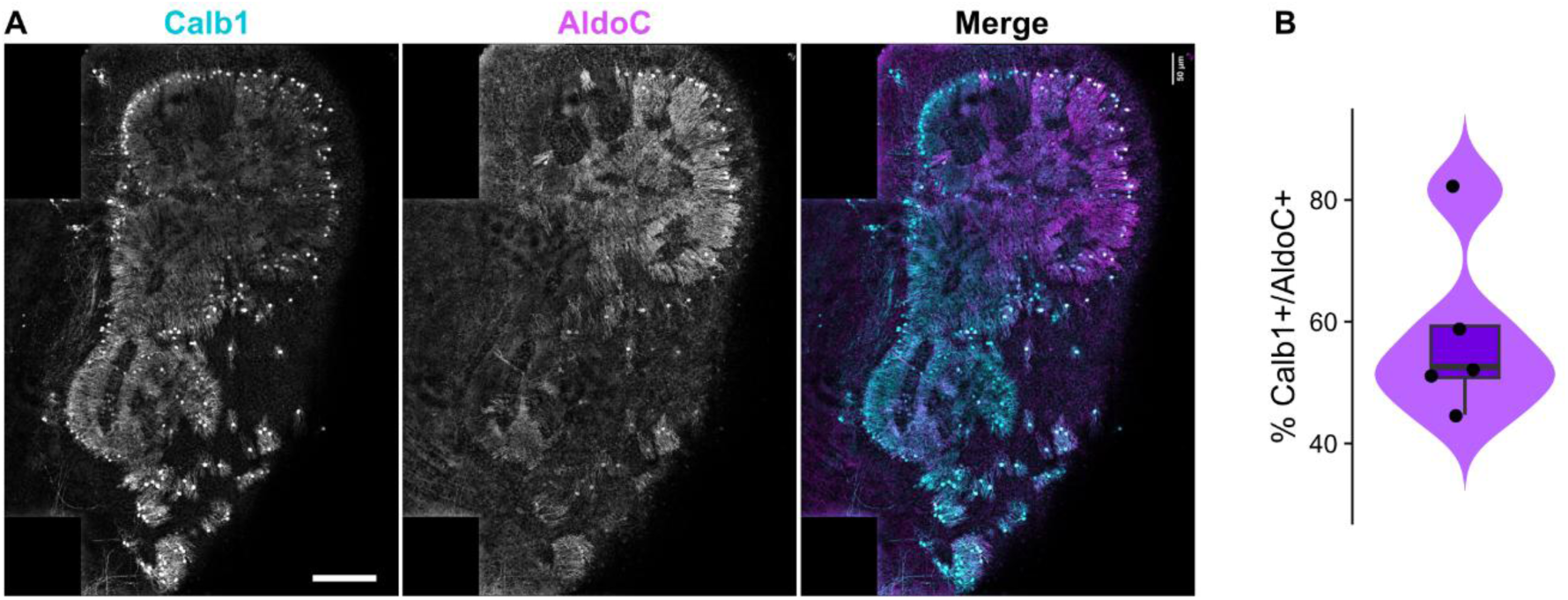
Zebrin-positive areas in hindbrain explants PCs. **(A)** Confocal mosaics (MIP) of a cerebellar plate immunostained for Calb1 (left, cyan in merge) and AldoC (middle, magenta in merge). Scale bar = 50 um. **(B)** Percentage of Calb1+/AldoC+ PCs per cerebellar plate.

**Supplementary Figure 3:**
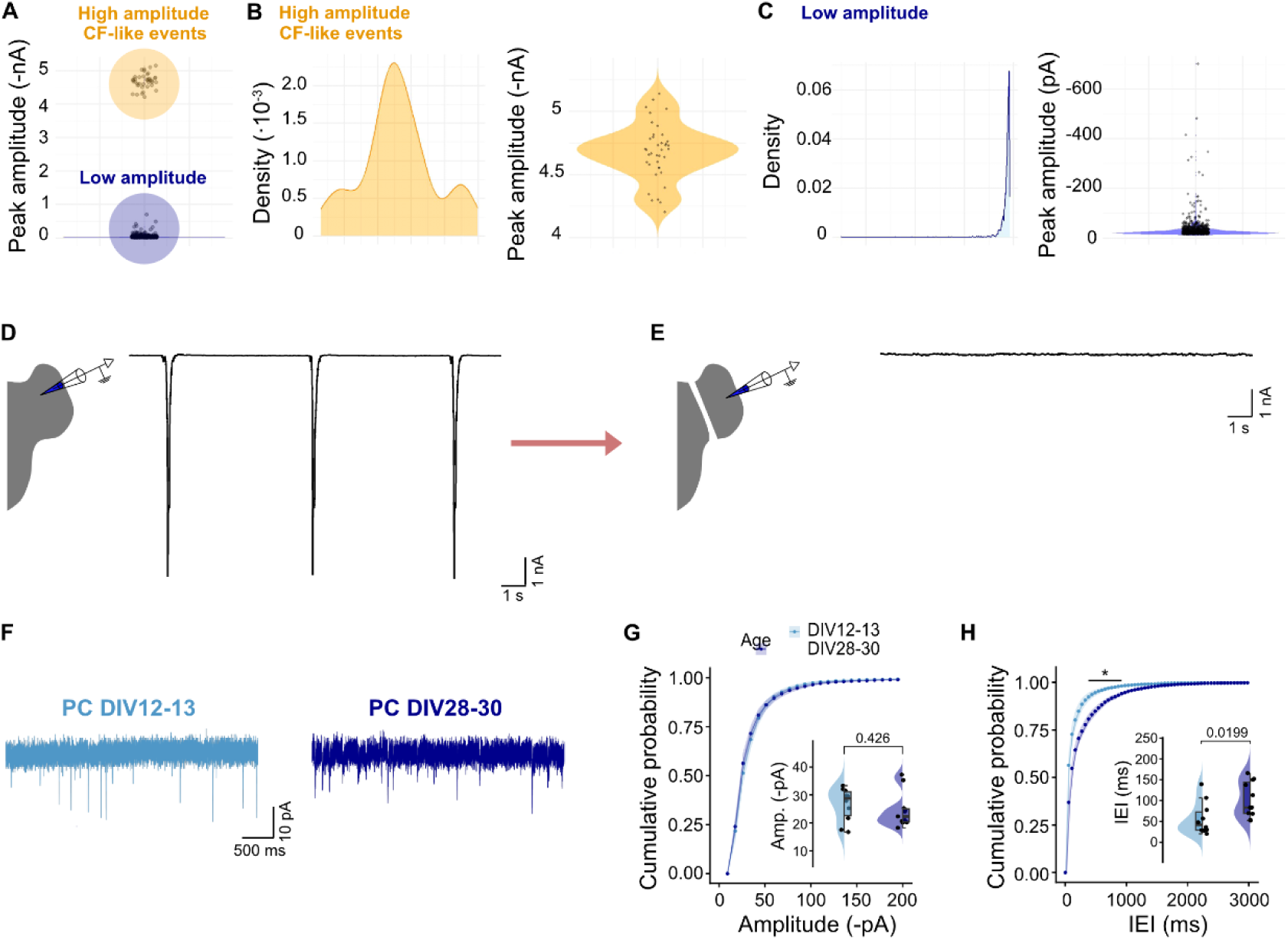
Electrophysiology recordings from PCs. (A-C) Events sorting from continuous voltage clamp PCs recordings for amplitude and frequency analysis. **(A)** Distribution of amplitudes for all the events detected in an individual PC (2’ recording). Events were separated into two amplitude-based categories: high (CF-like amplitude range) and low. **(B)** Density distribution (left) and range of amplitude values of “high amplitude” events. **(C)** Density distribution and range of amplitude values of “low amplitude” events. **(D)** Example trace of high amplitude events. **(E)** Example trace recorded from the same explant than (D) after acute dissection of brainstem. **(D-E)** Holding voltage = -70 mV. Scale bars correspond to 500 pA and 1 ms in both. **(F)** Representative traces of low-amplitude events, present in the inter-event interval of CF-like events (*Fig. 4G*). Scale bars correspond to 50 ms and 10 mV. **(G)** Cumulative distribution of low-amplitude sEPSCs. Median (IQR) were -28.7 (8.55) pA at DIV12-13 and -22.3 (4.63) pA at DIV28-30. **(H)** Cumulative distribution of IEI (ms) of low-amplitude sEPSCs. Median (IQR) IEI were 42.2 (43.2) ms at DIV12–13 and 83.4 (74.6) ms at DIV28–30. **(G-H)** Ribbon corresponds to S.E.M. Significance was determined using pointwise Wilcoxon rank-sum tests with Benjamini–Hochberg correction (FDR). P* < 0.05.

**Supplementary Figure 4:**
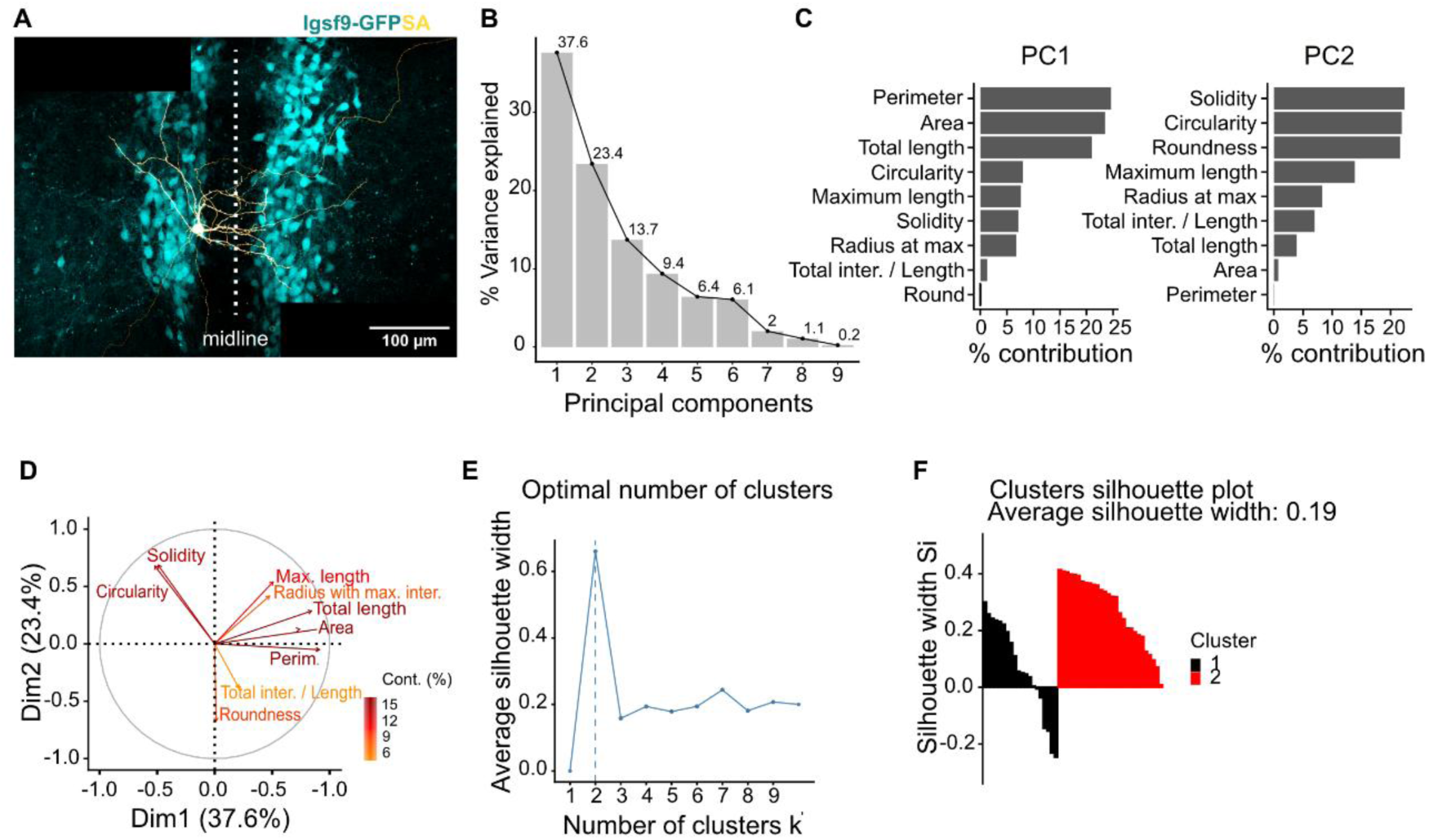
PCA and k-means evaluation. **(A)** Confocal image (MIP) of DIV13 biocytin-filled ION with its axon crossing to the contralateral brainstem. **(B)** Percentage of explained variance per principal component. **(C)** Percentage of contribution per variable to each principal component. **(D)** Contributing variables to principal components 1 and 2 (Dim1, Dim2) from cell shape descriptors and Sholl analysis. Arrows represent the direction and magnitude of each variable’s contribution to PCA space. **(E)** Average silhouette width calculated for the k-mean clustering obtained with k values ranging from 1 to 10. Peak is identified at k=2. **(F)** Silhouette plots showing the quality of k-means clustering. Each bar represents a cell and silhouette width reflects similarity to the assigned cluster relative to the nearest neighbouring cluster. Average silhouette width was 0.07 and 0.28 for cluster 1 and 2, respectively.

**Supplementary Figure 5:**
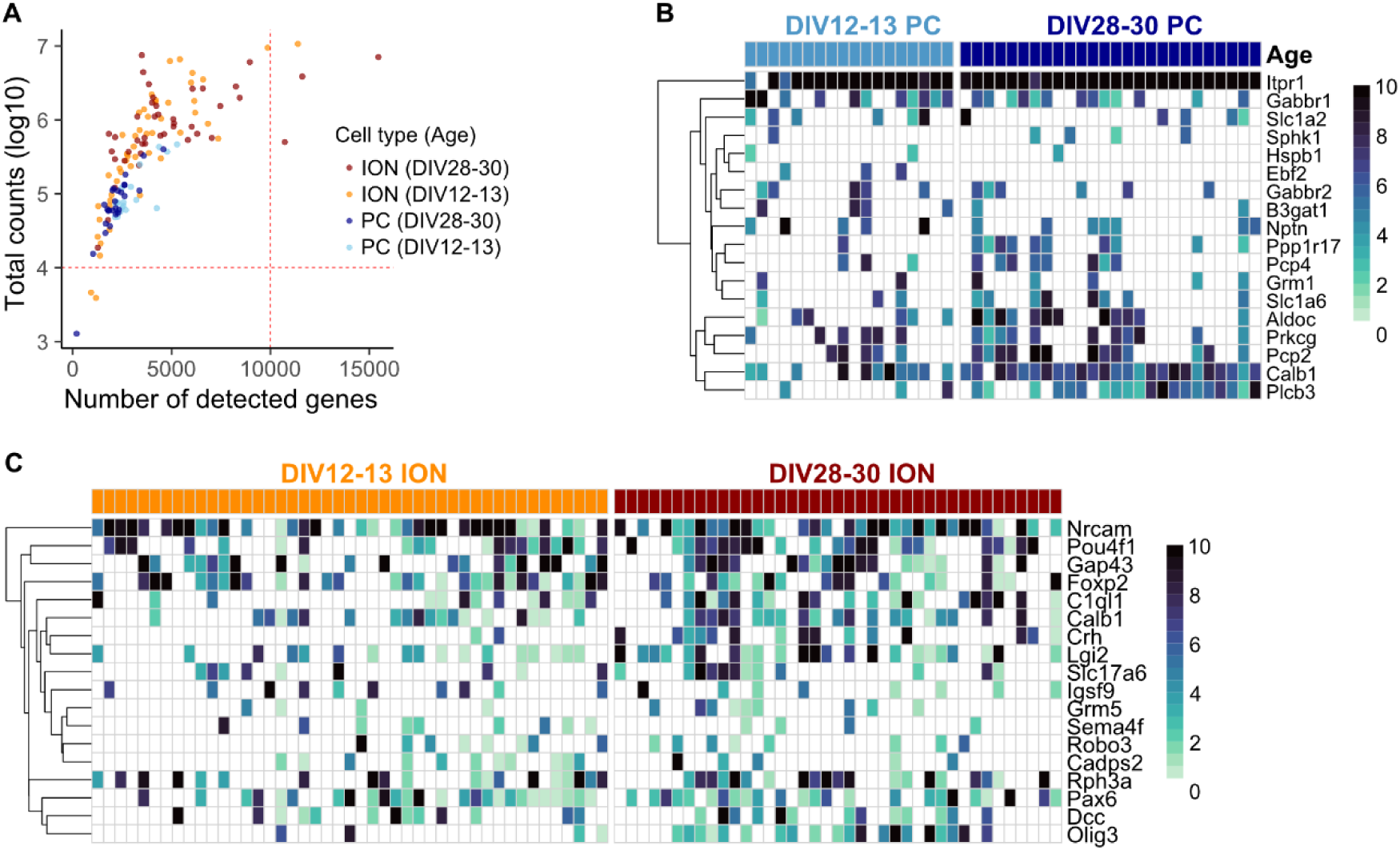
PatchSeq quality controls and marker genes expression. **(A)** Log10 of total counts sums and total number of detected genes per cell. Dashed red lines corresponds to QC thresholds. **(B)** Heatmap showing PC marker genes expression. **(C)** Heatmap showing ION marker genes expression. **(B-C)** Color scale corresponds to the log-normalized and min-max scaled counts.

